# Protocol for primary human lung organoid-derived air-liquid interface *in vitro* model to study response to SARS-CoV-2

**DOI:** 10.1101/2023.09.10.557067

**Authors:** Diana Cadena Castaneda, Sonia Jangra, Marina Yurieva, Jan Martinek, Megan Callender, Matthew Coxe, Angela Choi, Juan García-Bernalt Diego, Te-Chia Wu, Florentina Marches, Damien Chaussabel, Adolfo García-Sastre, Michael Schotsaert, Adam Williams, Karolina Palucka

## Abstract

This article presents a comprehensive protocol for establishing primary human lung organoid-derived air-liquid interface (ALI) cultures from cryopreserved human lung tissue. These cultures serve as a physiologically relevant model to study human airway epithelium *in vitro*. The protocol encompasses lung tissue cryostorage, tissue dissociation, lung epithelial organoid generation, and ALI culture differentiation. It also demonstrates SARS-CoV-2 infection in these cultures as an example of their utility. Quality control steps, ALI characterization, and technical readouts for monitoring virus response are included in the study.

For additional details on the use and execution of this protocol, please refer to Diana Cadena Castaneda et al (https://doi.org/10.1016/j.isci.2023.107374).

**Highlights:**

- Human lung tissue dissection, embedding in OCT blocks, and tissue cryopreservation.
- Thawing & lung tissue dissociation for lung epithelium organoid generation.
- Organoid-derived air-liquid-interface cultures for the study of viral infection.
- Bulk RNA-Seq, flow cytometry, viral titer, and imaging to follow response to virus.

**Graphical abstract:** 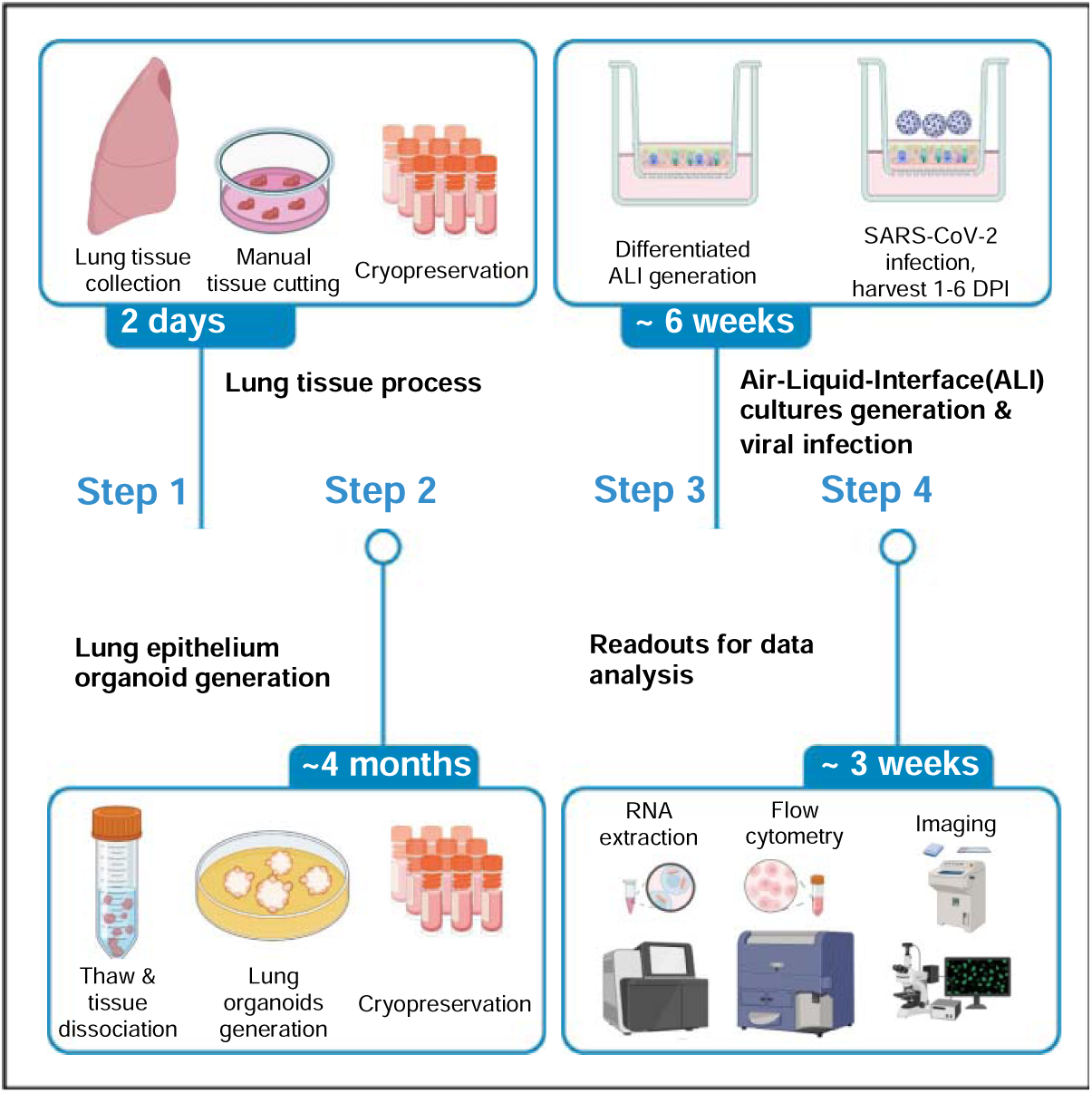

## Before you begin

The protocol below describes the generation of primary human lung epithelial organoid-derived air-liquid-interface cultures from lung tissue, and their use to investigate the impact and dynamics of infection with different SARS-CoV-2 variants. These cultures mimic in vivo airway epithelium architecture^2,3^ and are extensively used to study response to respiratory virus. The generation of lung epithelial organoids is based on methodology from Sachs et al^4^. and will be summarized here. The study involves transcriptional analysis (Bulk RNA), flow cytometry (dissociated ALI), viral titer measurements (apical supernatant washes), and imaging to assess the outcomes. All steps must be performed within a type II biological safety cabinet in a biological safety level 2 laboratory (BSL2). All work with SARS-CoV-2 variants must be performed under BSL3 conditions.

## Institutional permissions

De-identified human lung tissues were obtained from NDRI (Project: RPAK1 01), in compliance with relevant American laws, institutional and NIH/NIAID guidelines. Study procedures were performed in the context of U19AI142733 grant at the Jackson Laboratory. Before performing these procedures, permissions must be obtained from the relevant institutions.

## Human lung tissue processing & cryopreservation

Timing: 1-2 days.

On receipt of lung tissue, it is immediately processed. Throughout the process, attention is given to the lung tissue’s anatomical structure, facilitating dissection based on alveolar and bronchial areas. A portion of the lung is sectioned into approximately 3 cm x 3 cm pieces for embedding in OCT and snap frozen in liquid nitrogen. The remaining tissue is minced into smaller pieces, which are then cryopreserved in FBS with 10% DMSO and stored in liquid nitrogen before further processing. Around 25 of these small pieces are placed in each cryovial.

Following this procedure ensures the appropriate preservation of human lung tissue for organoid generation or other applications.

## Human lung tissue viable freeze thawing & lung airway organoid generation

Timing: Approximately 4 months

In brief, cryopreserved lung fragments (from 2-3 cryovials) are thawed and subjected to a dissociation process to yield a single-cell suspension, as outlined in the step-by-step section. The resultant single-cell suspension is mixed with cold Cultrex growth factor reduced BME type 2 (Matrigel-like matrix) and dispensed as droplets into a P24 well plate (approximately 3×10^5^ cells in 40 µl per well). These plates are then placed in a 37°C cell incubator for 20 minutes until gelation occurs. Following this, the wells are filled with complete media for organoids (AO), and the media is replenished daily. Organoids are passaged every 2 weeks until all connective tissue is eliminated, typically after 4-7 passages. This progression leads to the development of well-defined, spherical organoids devoid of connective tissue remnants. At this stage organoids are disrupted into single cells and cryopreserved (approximately 1.5×10^6^ cells/ml per cryovial) in FBS with 10% DMSO, and then stored in liquid nitrogen before subsequent processing to generate air-liquid interface (ALI) cultures.

**Critical**: Note that during all organoid-related steps, pre-coat tips, pipettes, and tubes with 5% BSA in to avoid organoid adherence to plasticware. Quality control via immunofluorescence staining is essential after 4-7 passages, once organoids are clear of connective tissue.

## Primary human lung organoid-derived air-liquid interface (ALI) culture generation, differentiation & viral exposure

Timing: Approximately 6 weeks

Cryopreserved single cells derived from human lung organoids (from 1 cryovial) are thawed and utilized for air-liquid interface (ALI) generation and differentiation. From 1 cryovial, 48 ALI cultures can typically be generated (P24 well plate format; one ALI culture = one insert seeded with 3×10^4^ cells per insert). This process comprises three main steps, as detailed in the step-by-step section: (1) cell expansion in submerged culture to obtain confluence at 100%; (2) initial differentiation in submerged cultures to foster tight junctions and barrier integrity, monitored by TEER values (>500 ohms.cm2). (3) Once TEER goals are achieved, cultures are transitioned to airlift by removal of apical media, which initiates the final differentiation into pseudo-stratified epithelia. Cultures are monitored for cilia beating and mucus production for a minimum of 4 weeks using a brightfield microscope. The resulting fully differentiated ALI cultures are ready for viral study protocols such as our investigation into SARS-CoV-2 response to variants (USA/WA1-2020, Wuhan-like), Beta (B.1.351), Delta (B.1.617.2), and Omicron (B.1.1.529, BA.1). Preceding viral exposure, apical mucus is removed by iterative washing with pre-warmed 1X PBS (37°C) On day 0 of viral exposure, 10^5^ PFU virus (or virus-free media for control conditions) is added apically. From days 1 to 6 post-infection, ALI cultures (each insert representing one culture) are harvested daily for kinetic experiments.

**Critical:** TEER evaluation is critical during ALI generation to monitor the health of the cultures. For viral exposure, use at least 25 µl or maximum of 100 µl of viral suspension for apical infection. Using larger volumes can induce tissue damage. Multiple readouts can be used to assess the response to the virus (see sections related to “Examples of readouts to assess response to viral exposure”).

## Examples of readouts to assess response to viral exposure

Timing: Approximately 3 weeks

We provide a succinct overview of potential readouts that could be employed to assess the response to viral exposure. Some of these readouts require a BSL3 setting, including flow cytometry, ALI dissociation & RNA extraction, and plaque assays for viral titers on the apical supernatant washes. However, other methods such as imaging (immunofluorescence, IF) or

GeoMX DSP on ALI frozen sections (following methanol free PFA 4% fixation overnight at room temperature (RT) and OCT embedding for snap-frozen tissue in LN) or dissociated ALI after Trizol treatment (for RNA preparation for sequencing) can be carried out outside a BSL3 environment following fixation or samples processing. However, all work must comply with institutional guidelines.

At each timepoint, SARS-CoV-2 infection can be assessed using flow cytometry on single-cell suspensions from dissociated tissue, tracking the expression of the viral nuclear protein (NP). Furthermore, apical washes can be collected to evaluate viral titer through plaque assays, providing insight into viral particle release on the apical side. Histocytometry, based on immunofluorescent image analysis, can also be used to evaluate infection through analysis of NP expression. Additionally, cellular ALI composition and relevant markers induced or elevated in response to the virus, such as CSF3 and CCL20, can be assessed through imaging (immunofluorescence, IF).

For transcriptional response to the virus, dissociated ALI cultures can be used for total RNA isolation using Direct-zol RNA MicroPrep kits, enabling sequencing and subsequent bulk RNA analysis. Moreover, NanoString’s GeoMx® Digital Spatial Profiler (DSP) can be performed on ALI culture frozen sections, offering insights into *in situ* RNA data in response to the virus.

**Critical:** Correct embedding of ALI cultures in OCT is crucial to obtain high-quality frozen sections. During immunofluorescence staining, it is essential to carefully choose conjugated and unconjugated antibodies with the correct secondary antibodies with appropriate fluorophores to avoid misleading staining.

The material and equipment section provides comprehensive recipes for all solutions, while the step-by-step section outlines the detailed procedures mentioned earlier. Solutions can be prepared beforehand, except for cell culture solutions which should be prepared fresh. For a comprehensive list of materials and equipment, please refer to the key resources table.

## Key resources table

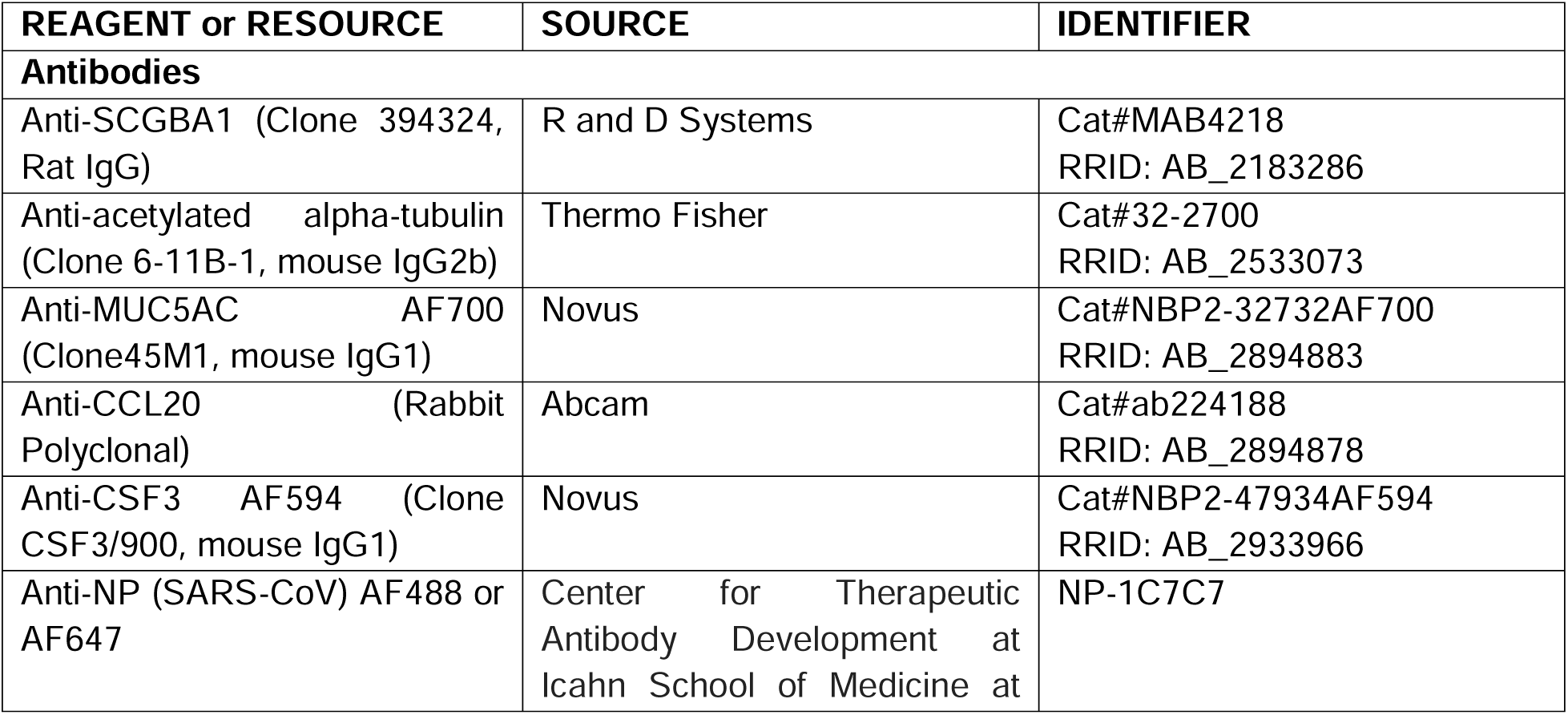
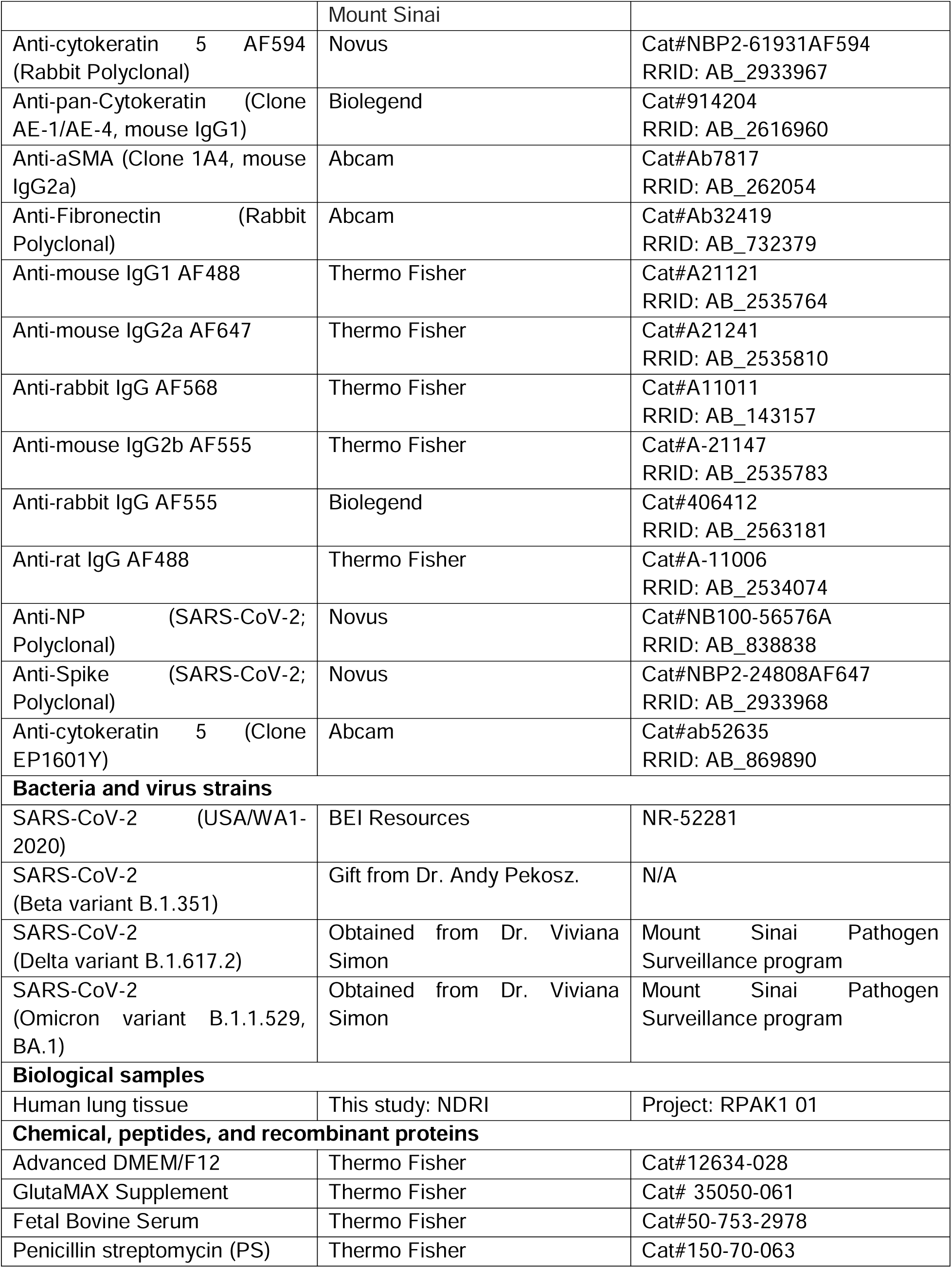
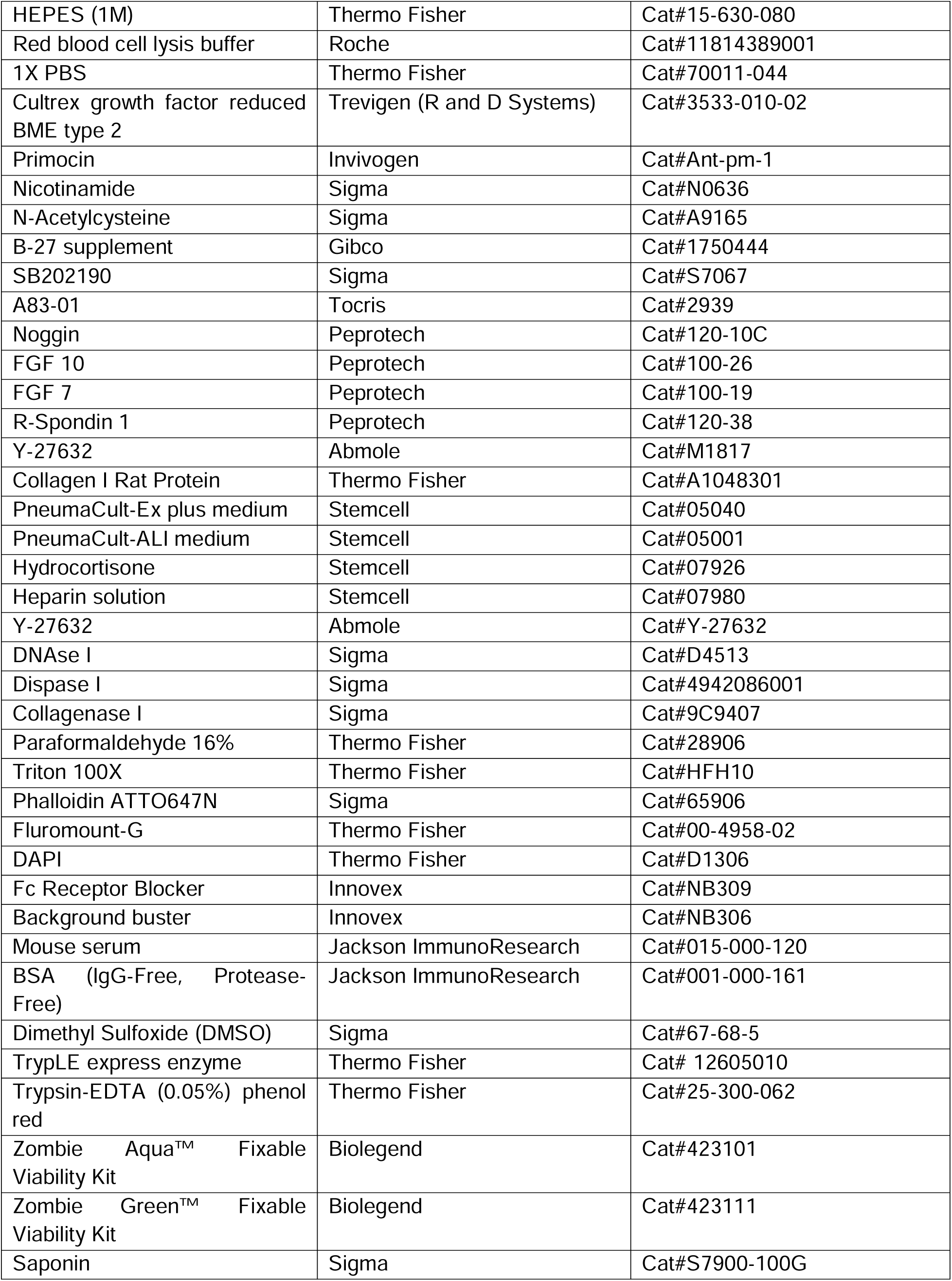
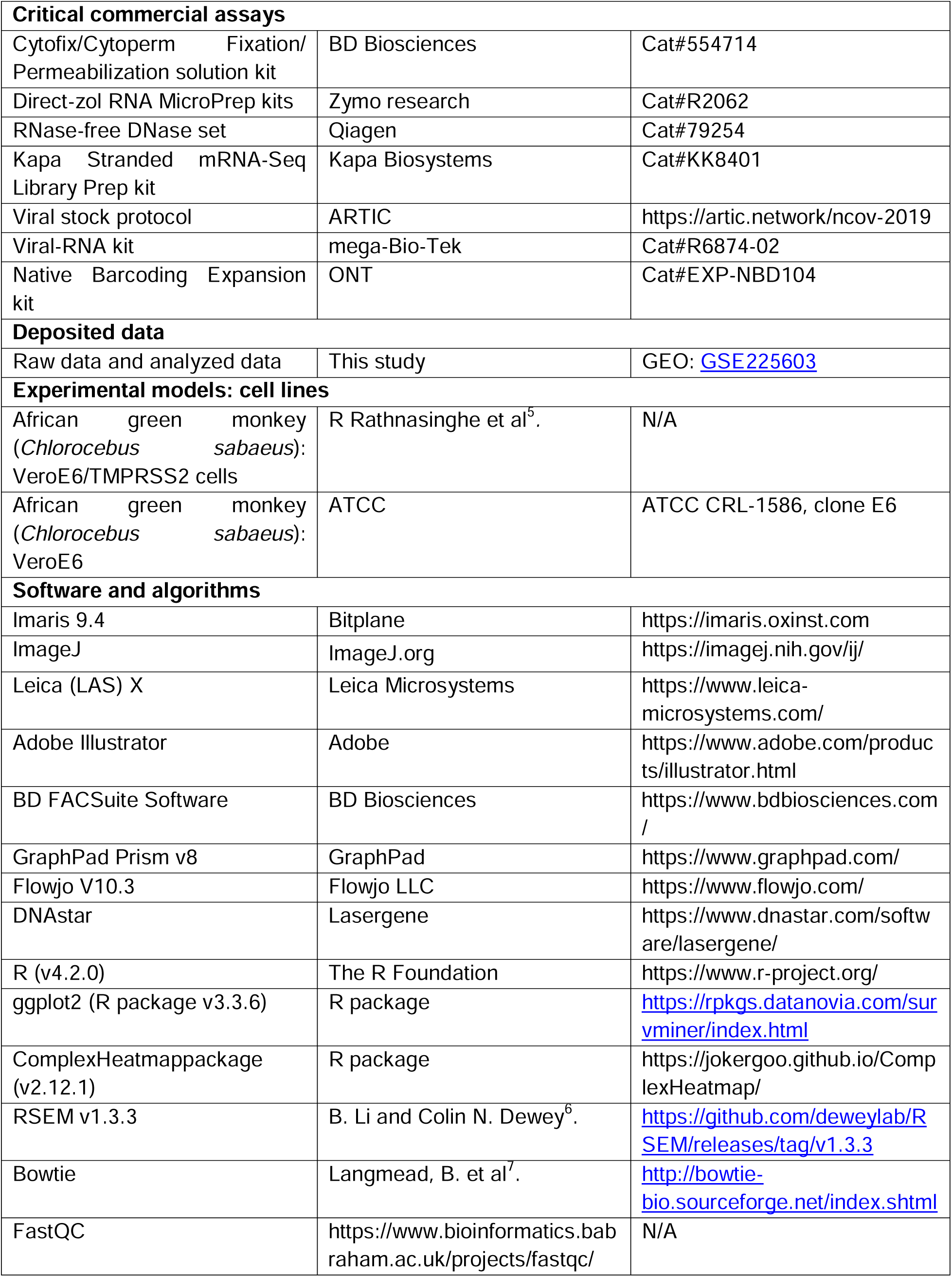
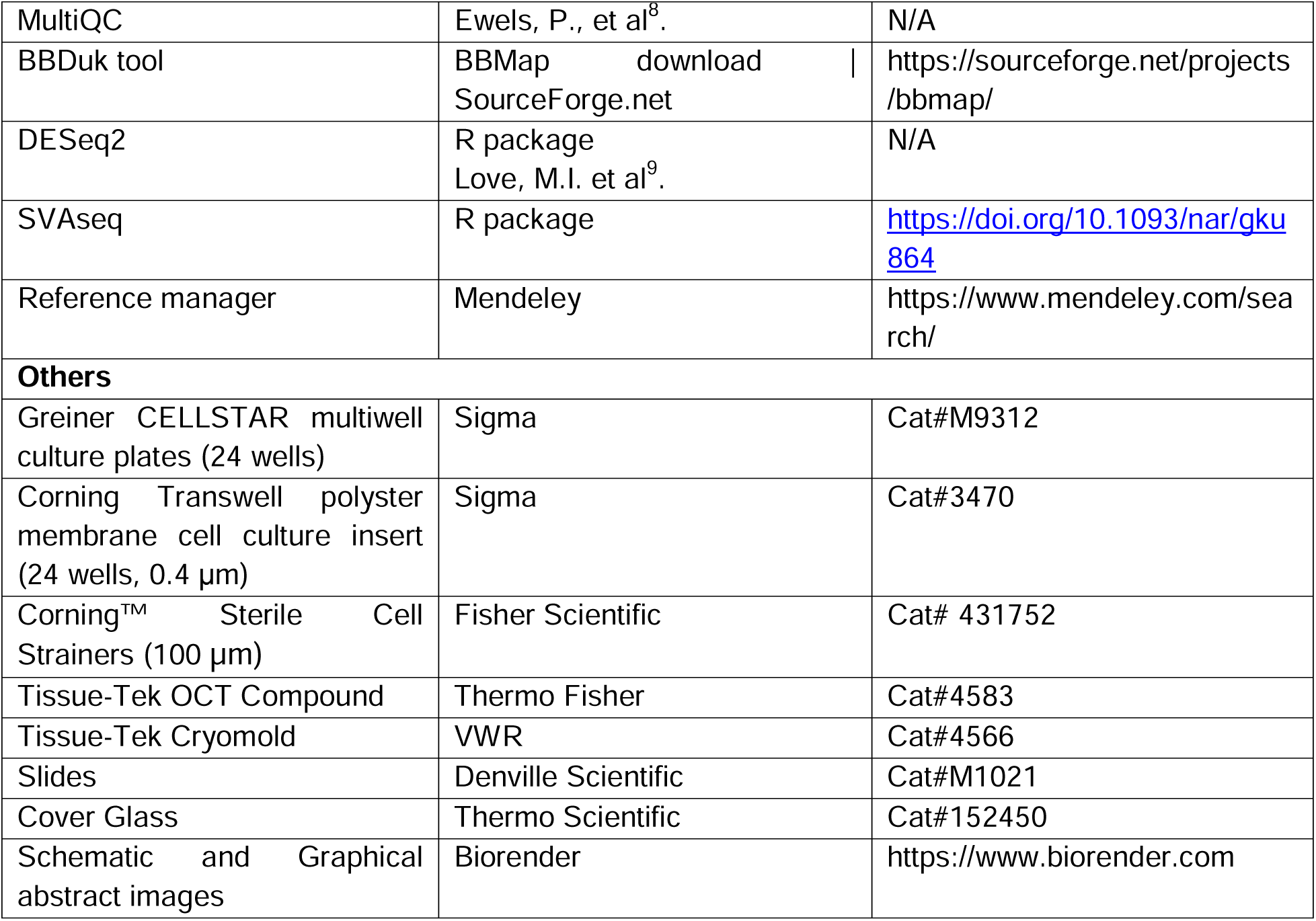

## Materials and equipment

### Human lung tissue processing & cryopreservation (1-2 days)

Equipment setup:

- Sterile material (autoclave): beakers (1000 ml in size), large Pyrex container (24 cm x 36 cm x 6 cm) for tissue reception, saran wrap to seal beakers, surgical blue wrap to avoid blood spills. Sterile scalpel, surgical forceps, fine-point forceps, surgical scissors.
- Pipettes: 1ml, 10ml, 25 ml.
- Petri dishes 100 x 15 mm (Thermo Scientific, 150350).
- Petri dishes 60 x 15 mm (Thermo Scientific, 174888).
- 50 ml centrifuge tubes, (Fisher Scientific, 339652).
- 2 ml cryovial tubes (Fisher Scientific, 431386).
- Tissue-Tek OCT Compound (Thermo Scientific, 4583).
- Tissue-Tek OCT cryomold (VWR, 4566).
- Optimized freezing container, Mr Frosty (Thermo Scientific, 5100-0001)
- Media for tissue preservation: 1XPBS (Thermo Scientific, 70011-044) supplemented with 50% FBS (Thermo Scientific, 50-753-2978).
- Media for viable freeze tissue cryopreservation: cold 90% FBS with 10% DMSO (Sigma, 67-68-5).

### Human lung tissue viable freeze thawing & lung airway organoid generation (∼ 4 months)

Equipment setup:

- Pipettes: 1ml, 10ml, 25 ml.
- Petri dishes 60 x 15 mm.
- 50 ml centrifuge tubes.
- Tube rotator (Miltenyi Biotec, MACSmix™ Tube Rotator, 130-090-753).
- Sterile slides (Globe Scientific, #1324W).
- Sterile Cell Strainers, 100 µm (Fisher Scientific, 431752). It is important to pre-wet strainers before filtering tissue or cells suspensions with 1-2 ml 1X PBS.
- Piston from insulin syringe (BD, 324921).
- Cultrex growth factor reduced BME type 2 (R and D, 3533-010-02). It is important to melt this Matrigel-like matrix reagent 24hr prior to viable frozen tissue processing in the fridge at 4°C.
- Greiner-Cellstar 24 well culture plates (Sigma, M9312).
- TrypLE express enzyme (Thermo Fisher, 12605010)

Preparation of all media to be used during viable freeze thawing and lung airway organoid generation are described here below.

Preparation of basal media for airway organoid (AdDF+): this media can be used as basal media for tissue digestion media or for complete media for airway organoid media (AO).

| Reagent | Stock concentration | Final concentration | Storage |
| --- | --- | --- | --- |
| Advanced DMEM/F12 | Ready to use | N/A | 4°C for 1 year |
| Penicillin/Streptomycin | 100X | 1X | -20°C for 1 year |
| Hepes | 1M | 10mM | 4°C for 1 year |
| GlutaMAX Supplement | 100X | 1X | 4°C for 1 year |

Preparation of media for tissue digestion: AdDF+ media supplemented with 1-2mg of I collagenase (Sigma-C9407, stock concentration: 200mg/ml, −20°C for 1 year).

Preparation of complete media for airway organoid generation (AO): this media is used for airway organoid selection, formation, and maintenance. Important to aliquot every growth factor or chemical regent to avoid freeze and thaw cycles.

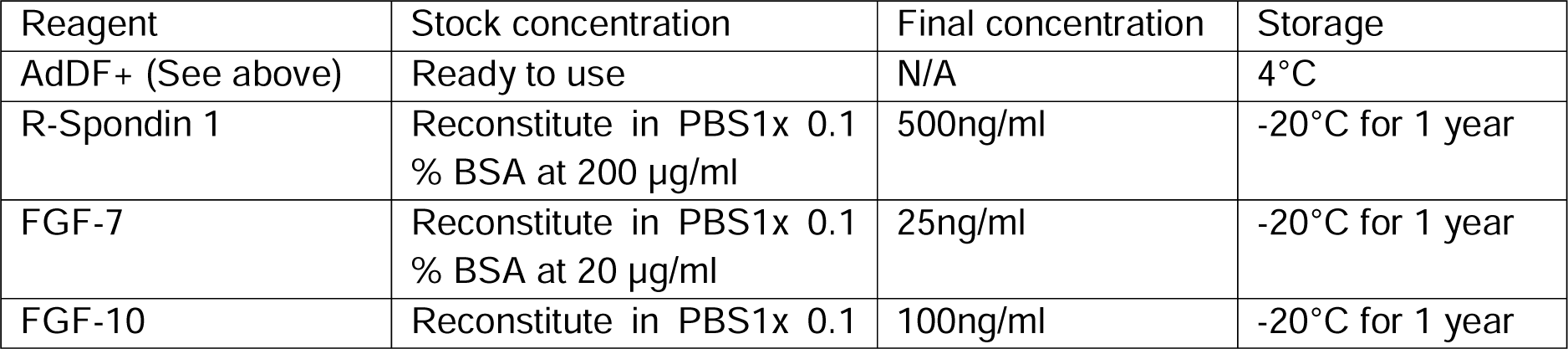
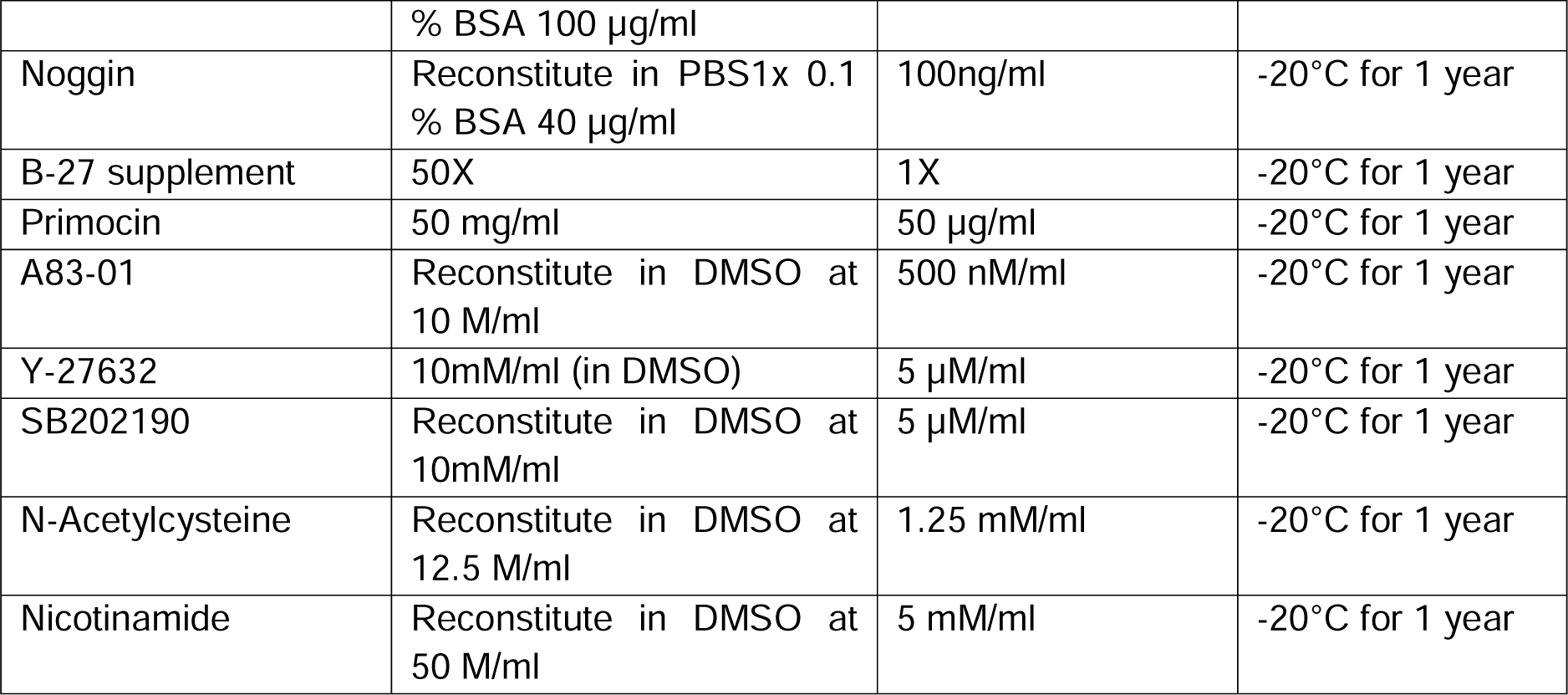

Preparation of media to stop tissue digestion reaction: 1X PBS supplemented with 2% FBS.

### Primary human lung organoid-derived air-liquid interface (ALI) generation, differentiation & viral exposure (6 weeks)

Equipment setup:

- Pipettes: 1ml, 10ml, 25 ml.
- 50 ml centrifuge tubes.
- EVOM^2^ ohm meter (WPI). For more details on how to use the device refer directly to <u>WPI supplier website</u>.
- Corning Transwell polyster membrane cell culture insert (24 wells, 0.4 μm; Sigma, 3470).
- Collagen I Rat Protein, Tail, at a final concentration of 30µg/ml in 1X PBS (Thermo Fisher, A1048301).
- Sterile forceps.
- StemCell Pneumacult Ex-Plus (#05040), and StemCell Pneumacult ALI Maintenance media (05001).

Preparation of all media to be used during ALI generation and viral exposure are described below. This process comprises three main steps (1-3):

Preparation of complete media for ALI culture expansion (Step 1): this media is required on apical (100 μl) and basal side (500 μl) after cell seeding for the first 10-12 days, depending on the donor.

| Reagent | Volume | Final concentration | Storage |
| --- | --- | --- | --- |
| PneumaCult™-Ex Plus Basal Medium | 490 ml | N/A | 4°C for 1 year |
| PneumaCult™-Ex Plus 50X Supplement | 10 ml | 1X | -20°C for 1 year |
| Hydrocortisone Stock Solution | 500 µl | 1X | -20°C for 1 year |
| Penicillin/Streptomycin | 5 ml | 1X | -20°C for 1 year |
| Y-27632 (do not add to entire bottle, add fresh to aliquot each day) | For 50 ml add 50 µl | 0.01 mM/ml (stock at 10 mM/ml) | -20°C for 1 year |

Preparation and usage of complete media for ALI culture differentiation and maintenance:

- Confluence and Initial Media Change (Step 2): After achieving full confluence of ALI cultures (around 10-12 days post-seeding), change the media on both the apical and basal sides. Use media designed for ALI culture differentiation and maintenance, enriched with Y-27632. This compound is essential for the initial phase of pseudo-stratified epithelium differentiation, spanning approximately 5-7 days (varies based on donor characteristics).
- TEER Goal Achievement and Transition to Air-Lift (Step 3): Once the targeted TEER values surpass 500 ohms.cm^2^, commence the air-lift phase. At this point, discontinue the addition of media to the apical side while maintaining media supply to the basal side. When ALIs are air-lifted, supplementation with Y-27632 should be halted. This marks the transition to the second phase of differentiation, lasting approximately 4 weeks.

By supplementing with Y-27632 and altering media application according to the differentiation phases, you facilitate the progression of ALI cultures towards a pseudo-stratified epithelial state.

Basal media for ALI culture differentiation and maintenance

| Reagent | Volume | Final concentration | Applications |
| --- | --- | --- | --- |
| PneumaCult-ALI Basal Medium<br>(Store at -20°C for 1 year) | 450 ml | N/A | Prepare, aliquot, & store at -20°C (for 1 year), thaw when ready to use. |
| PneumaCult-ALI 10X Supplement<br>(Store at -20°C for 1 year) | 50 ml | 1X |  |
| Penicillin/Streptomycin | 5 ml | 1X |  |

Complete media for ALI culture differentiation and maintenance

| Reagent | Volume | Final concentration | Applications |
| --- | --- | --- | --- |
| Basal media for ALI culture differentiation and maintenance (See above) | 10 ml | 1X | Aliquot stored at -20°C (for 1 year), thaw when ready to use. |
| Hydrocortisone Stock Solution<br>(Store at -20°C for 1 year) | 200X (50µL/10mL Complete Base) | 1X | Add only desired amount when using media (add to the thawed aliquots of PneumaCult ALI Complete Base) |
| Heparine Solution<br>(Store at 4°C for 1 year) | 500X (20µL/10mL Complete Base) | 1X |  |
| PneumaCult-ALI Maintenance Supplement (Store at -20°C for 1 year) | 100X (100µL/10mL Complete Base) | 1X |  |
| Y-27632<br>Add only desired amount (do not add to entire bottle)<br>(Store at -20°C for 1 year) | For 10 ml add 10 µl | 0.01 mM/ml (stock at 10 mM/ml) | Include Y-27632 to media prior to airlift, do not include afterward. |

All experiments involving live SARS-CoV-2 should be conducted in a biosafety level 3 (BSL-3) facility Ensure that all necessary safety measures are followed during the viral exposure experiments.

- Media to apply on basal side of ALI culture during viral exposure experiments: PneumaCult-ALI Basal Medium supplemented with PneumaCult-ALI 10X supplement (1X final) and Penicillin/Streptomycin (1X final) is used directly without any other supplement.
- SARS-CoV-2 USA/WA1-2020 virus or variant (Beta; Delta; Omicron) to be used individually at 10^5^ PFU final/well.
- All viral stocks should be sequenced to confirm genomic integrity according to the ARTIC protocol as previously described (https://artic.network/ncov-2019)^5^. Aliquots should be stored in a secured −80L°C freezer until use.

### Example of readouts to assess response to viral exposure (∼ 3 weeks)

- Preparation of material and reagents for plaque assays.

At all time-points viral supernatants are collected by adding 150μl of 1X PBS on the apical side and incubating for 15 min at 37°C. Then, supernatants should be stored at −80°C until processed. All experiments involving live SARS-CoV-2 isolates should be conducted in a biosafety level 3 (BSL-3) facility.

Equipment setup:

- Pipettes: 1ml, 10ml, 25 ml.
- 1.5 ml Eppendorf tubes.
- 1% Bovine pre-seeded VeroE6 cell (for USA/WA1-2020) or Vero-E6 TMPRSS2 (for Beta, Delta and Omicron) monolayers.
- 2% Agar (w/v): dissolve 2g of purified Oxoid (LP0028) agar in 100 ml water.
- Prepare 2X MEM media: 200ml of 10X MEM supplemented with 20ml of 200mM L-glutamine, 32 ml of 7.5% sodium bicarbonate, 20ml of 1M HEPES, 20 ml of 10mg/ml Pen/Strep, 12ml of 3.5% BSA and 696 of water. Filter sterilizes using a Millipore (0.22 μM) vacuum-driven filter.
- 1% DEAE Dextran (w/v): dissolve 0.5g of DEAE Dextran in 50 ml of water. Filter sterilizes using a Millipore (0.22 μM) vacuum-driven filter.
- 2% Oxoid agar mixed with 2X MEM solution supplemented with DEAE Dextran and TPCK/trypsin. Example for 25 ml in a conical tube (50 ml): 4ml of water, 250 μl of DEAE Dextran, 12.5 ml of 2X MEM, 25 μl of TPCK/trypsin (stock 1mg/ml, working 1μg/ml), this mix is kept at 37°C. When ready to add overlay to cells, combine the 8.5ml agar to the mixture in conical tube and immediately add.
- PFA 4% fixation for immune staining (Thermo Fischer, 28906).
- To reveal viral infection: Anti-SARS-CoV-2 NP antibody (1C7C7), and HRP-conjugated secondary anti-mouse antibody (1:5000).
- True Blue peroxidase substrate (Sera Care).

- Preparation of reagents for ALI dissociation, intracellular staining for infection (Flow cytometry) and RNA extraction.

At all time-points, Infected and uninfected ALI are dissociated: separately, one insert for flow cytometry and one insert for RNA extraction. See the step-by-step section for the detailed procedure. Those readouts should be conducted in a biosafety level 3 (BSL-3) facility.

Equipment setup:

- Pipettes: 1ml, 10ml, 25 ml.
- Trypsin-EDTA (0.05%) phenol red (Thermo Fisher, 25-300-062).
- Prepare a fresh Dispase I/DNase I solution in 1X PBS with a final concentration of 1.9U/ml diaspase I (Sigma, 4942086001) and 0,1mg/ml of DNase I (Sigma, D4513).
- Media to stop tissue digestion reaction: 1X PBS supplemented with 2% FBS.
- 1.5 ml Eppendorf tubes.
- Flow cytometry:

- Viability assessment: staining in 1X PBS with Zombie aqua (#423101) or zombie green (#423111); (Biolegend).
- Cytofix/Cytoperm Fixation/ Permeabilization solution kit (BD, 554714).
- SARS-CoV-2 N antibody (1C7C7, AF488 or AF647)
- Buffer for cell suspension: 1X PBS+ 1% BSA.
- BD FACSuite Software and Flowjo.
- RNA extraction: Extracted RNA should be stored at −80, until used.

- Direct-zol RNA MicroPrep kits (zymo research, R2062).
- RNase-free DNase set (Qiagen, 79254).
- PolyA mRNA selection and Kapa Stranded mRNA-Seq Library Prep kit (Kapa Biosystems, KK8401).
- DNAstar (Lasergene).
- Preparation of ALI cultures for imaging and Nanostring GeoMx whole transcriptome atlas (WTA).

Equipment setup:

- Pipettes: 1ml, 10ml, 25 ml.
- Tissue-Tek OCT Compound (Thermo Scientific, 4583).
- Tissue-Tek OCT cryomold (VWR, 4566).
- Cryostat Leica
- Immunofluorescence (IF):

- PFA 4% (methanol free) fixation for immune staining (Thermo Fischer, 28906).
- Cold acetone (−30°C). To be used in a fume hood.
- Permeabilization buffer: 1X PBS with 0.1% Triton 100X.
- Saturation (Innovex, NB309) and background reagents (Innovex, NB306).
- Staining buffer: 1XPBS with 5% BSA (IgG-Free, Protease-Free), (Jackson ImmunoResearch, 001-000-161) and 0.05% Saponin (Sigma, S7900-100G).
- Primary and secondary antibodies (see key resources table).
- Mouse serum, (Jackson ImmunoResearch, 015-000-120).
- Phalloidin ATTO647N, (Sigma, 65906).
- DAPI (Thermo Fisher, D1306)
- Hydrophobic pen (pap-pen, Daido Sangyo, N33)
- Mounting media (Thermo Fisher, 00-4958-02).
- Superfrost plus Slides (Denville Scientific, M1021) and cover-glass (Thermo Fisher, 152450)
- GeoMx WTA: Slides with 6µm thickness sections of ALI cultures at different conditions should be stored at −80°C until process.

- PFA 4% fixation for immune staining (Thermo Fischer, 28906).
- Tissue is prepared according to MAN-10131-03. For more details, please refers to <u>Nanostring website</u>.
- Primary antibodies: Nuclei Syto83 (Nanostring), CK5 (Abcam, ab52635, FITC), SARS-CoV-2 Spike (Novus, NBP2-24808AF647), and SARS-CoV-2 Nucleocapsid (Novus, NB100-56576).
- Secondary antibodies: Anti-rabbit IgG AF555 (Biolegend, 406412).
- Viral-RNA kit (mega-Bio-Tek, R6874-02)
- Native Barcoding Expansion kit (ONT, EXP-NBD104).
- DNAstar (Lasergene).

**Critical**: Reagents or biological materials may be harmful and/or toxic, therefore use appropriate laboratory equipment. For flow cytometry, IF and GeoMx staining make fresh reagents and antibody mix for each staining, keep on ice in the dark.

### Step-by-step method details

#### Human lung tissue processing & cryopreservation

Timing: 1-2 days.

This step allows the selection of suitable tissues for cryopreservation and airway organoid generation based on tissue anatomy, elimination of necrotic or damaged areas and cryopreservation of multiple vials from different regions of the lung.

1. Lung tissue processing (Figure 1): Receive human lung tissue under clean/sterile conditions.
  a. Transfer the lung tissue to a sterile beaker (1000 ml) containing 1X sterile PBS (400-500 ml to cover the tissue) supplemented with 50% FBS. Seal the beaker with sterile aluminum foil, then, parafilm and keep it at 4°C until tissue processing.
  b. Transfer the lung into a large sterile Pyrex container (24 cm x 36 cm x 6 cm) and anatomically orientate the lung tissue. For example, for left lung, place the lung facing the researcher in a way that distal area (alveolar part) is on the left and proximal part (bronchial part) in on the right close to the trachea, (See Figure 1, Step1).
  c. Cut lung tissue into slices (approximately 4-5 cm thick) with a sterile scalpel in function of anatomical orientation, starting from the apex (upper part) to bottom part of the lung, then from distal (more alveolar) to proximal (more bronchial) (See Figure 1, Step 2).
  d. Cut each lung tissue slice into smaller pieces (approximately 3 cm x 3 cm in size), with a sterile scalpel (See Figure 1, Step 3).
  e. Embed representative lung pieces from each region (approximately 3 cm x 3 cm in size) in OCT and snap freeze in liquid nitrogen. Name each OCT tissue block based on anatomical region and donor code: for example, LA for “alveolar” and LB for “bronchial” (See Figure 1, Step 4(a)).
  f. Mince the remaining tissue into smaller pieces, of approximately 3 mm x 3 mm in size first using a sterile scalpel then surgical scissors. Transfer these smaller lung tissue pieces into cryovials with around 20-25 pieces per vial (2ml cryovial size) with forceps. Immediately add 1 mL of cold cryopreservation media (FBS 10% DMSO) to each cryovial and transfer to an optimized freezing container and store at −80°C for 24hrs before transferring to liquid nitrogen for long-term storage. Similarly, label each cryovial based on anatomical region, donor code, and date: (See Figure 1, Step 4(b)).

**Figure 1:**
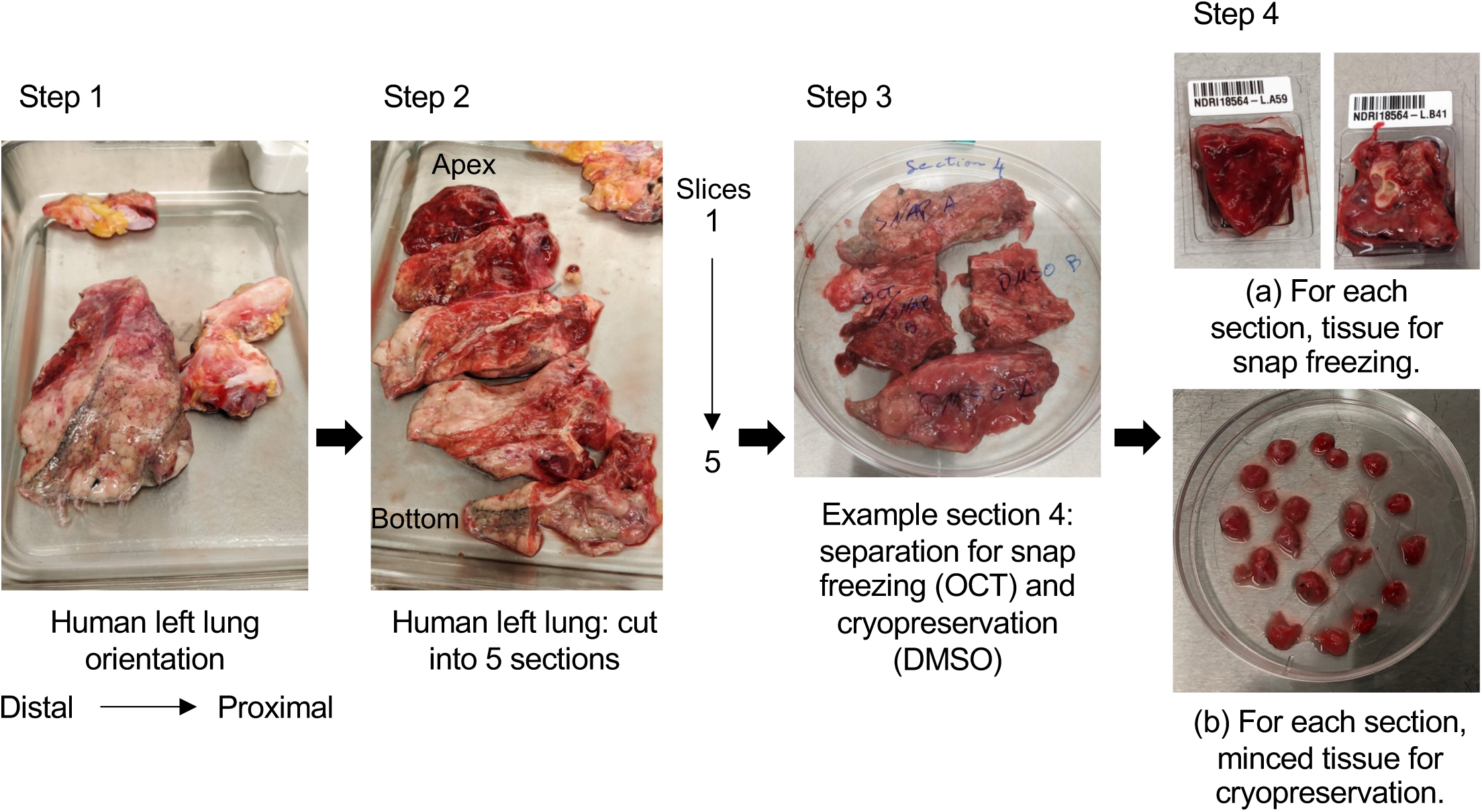
Set up for human lung tissue processing and cryopreservation: representative pictures for lung tissue processing. (Step 1) The left lung, including a partial portion of the trachea, is used for the processing. The tissue is oriented with the distal (alveolar) portion on the left and the proximal (bronchial) portion on the right. (Step 2) Cut the tissue into five sections from the apex (upper part) to the lower part. (Step 3-4) Cut each section cut into smaller pieces. Track the pieces from different areas of the lung as follows: for example, those from the alveolar region are named LA, and those from the bronchial region are named LB. From each region, either snap-freeze small pieces in OCT for histology or subject them to cryopreservation (10% FBS and DMSO).

**Critical**: By applying this method, the human lung tissue sections are processed and categorized based on anatomical region.

#### Human lung tissue viable freeze thawing & lung airway organoid generation

Timing: Approximately 4 months

This section describes the workflow to culture primary airway organoids, including the isolation, expansion, and cryostorage of lung organoid-derived progenitor cells, along with quality control screening of organoids for epithelial enrichment (Figure 2).

2 Processing of viable frozen lung pieces and digestion for airway organoid generation.
  a. Prepare basal media for organoids (AdDF+) which will be used for washings steps and complete media for organoids (AO) which will be used for tissue digestion and organoid generation. See Materials and equipment section for the media preparation. Media should be freshly prepared and pre-warmed (37°C) prior to tissue thawing. In addition, prepare media to stop tissue digestion (1X PBS supplemented with 2% FBS) and thaw Cultrex growth factor reduced BME type 2 (Matrigel-like matrix), 24hr prior to viable frozen lung pieces processing (at 4°C in the fridge).
  b. Thaw the cryopreserved tissue by placing 2-3 cryovials into a dry bath at 37°C for 5 min.
  c. Immediately, transfer the minced tissue into a 50 ml tube containing pre-warmed AdDF+ media (20-30 ml) and centrifuge for 5 min at 300 G (RT) to remove the DMSO.
  d. Resuspend the minced tissue in 10 ml of complete AO containing 1–2 mg ml collagenase I and incubate on an orbital shaker at 37°C for 1h to allow tissue digestion. <u>Troubleshooting 1.</u>
  e. Shear the digested tissue between sterile slides and strain through a 100µm filter. To improve the straining use the piston of an insulin syringe to press the tissue through the filter. For tissue remaining in the filter, repeat the shearing and filtration steps to remove as much connective tissue as possible to obtain a single cell suspension.
  f. Rinse the filter and stop the digestion process by adding 1X PBS supplemented with 2% FBS (RT). Then, centrifuge for 5 min at 300 G (RT). Resuspend the pellet in 1X PBS (approximately 5 ml).
  g. In case a visible red pellet is present, centrifuge for 5 min at 300 G (RT) and proceed to erythrocyte lysis using 2 ml red blood cell lysis buffer for 5 min at room temperature before the addition of 10 ml AdDF+. Strain cells through a 100µm filter using a new insulin syringe piston and centrifuge at 300 G (5min, RT). <u>Troubleshooting 2.</u> Resuspend the pellet in AO media (1 ml).
  h. Count viable cells before proceeding to organoid generation. At this point, we expect to obtain 5 to 7.5×10^6^ cells with a viability of 80-90%.
3 Primary human lung organoid generation is performed based on Sachs et al^4^ and will be described briefly (Figure 2A).
  a. Immediately, resuspend the lung cell suspension (7.5×10^6^ cells/ml final) in liquid Cultrex growth factor reduced BME type 2, (1 mg/ml final). Keep the suspension on ice until you are ready to proceed to cell culturing. Mix and dispense approximately 300 000 cells/40µl as a dome (1 dome per well) into a pre-warmed Cellstar 24 well culture plate. Flip the plate upside-down and incubate in a humidified 37°C/5% CO2 incubator at ambient O_2_ for 20 min to allow jellification. <u>Troubleshooting 3</u>.
  b. Flip the plate right side up and gently add 400 µl of pre-warmed AO complete media supplemented with factors allowing the enrichment in lung epithelia cells.
  c. Chang media every day, being careful not to disturb the Cultrex growth factor reduced BME type 2 dome. Passage organoids every 2 weeks as described below.
  d. To passage organoids, remove media, add cold 1X PBS and incubate the plate on ice for 30 min.
  e. Pipette (1 ml pipette) up/down to break the domes, combine all the wells and collect the organoids in a50 ml tube, centrifuge at 300 G for 5 min (7°C).
  f. To fully dissociate the organoids into single cells, add pre-warmed 1X TrypLE express, incubate 5 min at room temperature.
  g. Mechanically (by pipetting up/down) dissociate the organoids and stop the reaction by adding ice cold 1X PBS supplemented with 2% FBS and centrifugation at 300 G 5 min (7°C).
  h. Verify under the microscope that dissociation is complete. This should be evident as a distinct single cell suspension, free from intact organoids. If necessary, repeat steps from (e) to (h).
  i. Centrifuge (300G, 5 min, at RT) and resuspend the cells in Cultrex growth factor reduced BME type 2 repeating the steps starting from (a) to (c).
  j. After each dissociation, count cells and cryopreserve a proportion when possible (ideally at least 3-4 cryovials with 1.5×10^6^ cells/ml/vial) in FBS with 10% DMSO. Having these back-up stocks for later expansion is critical as bacterial or fungal contamination may occur during the expansion, which is a risk while working with primary tissues. Once the organoids are clear of the connective tissue, (after approximately 4-7 passages), generate a large frozen stock of dissociated organoids at 1.5×10^6^ cells/cryovial. At this stage, there are normally 3-4 24 wells plates of organoids in culture that would be expected to produce 20-25 cryovials.

**Figure 2:**
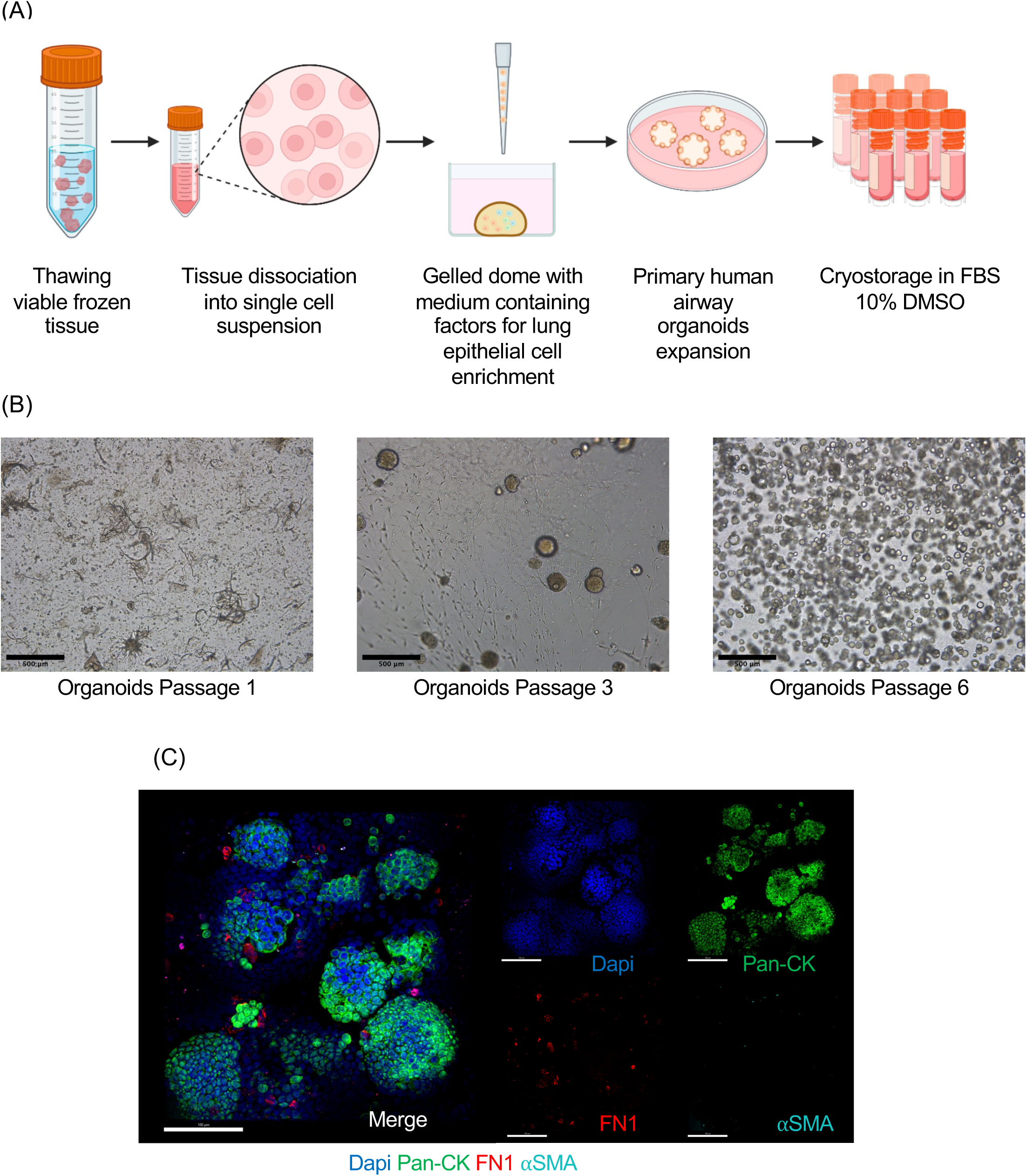
Viable frozen tissue processing and generation of primary airway lung organoids: (A) Schematic experimental design for primary human lung organoid generation: viable frozen tissue is thawed and digested to obtain a single cell lung suspension containing airway cell progenitors. The cell lung suspension is resuspended in Cultrex, dispensed as dome, and cultured in medium containing factors allowing airway organoid expansion and lung epithelium enrichment. After each passage dissociated organoids are cryopreserved. B) Representative photomicrographs of primary lung organoids at different passages captured using a brightfield microscope. Images were generated using ImageJ. Scale bars 500µm, in black on the left corner. (C) Representative immunofluorescent (IF) images of whole mounted lung organoids showing markers for epithelial cells (PAN-CK, green), extracellular matrix (Fibronectin, red), and nuclei (DAPI, blue). Pan-CK is present exclusively in epithelial cells and fibronectin allows visualization of the presence of remaining extracellular matrix from tissue dissociation. The fibronectin disappears after four passages and characterizes the enrichment of lung epithelial cells. Scale bar 100 µm, in white in the left corner.

**Critical**: For the first 1-4 passages, there is a lot of connective tissue that becomes progressively diluted with each passage (Figure 2B). Once spherical organoids are obtained, and domes are clear of connective tissue a quality control step is required (Figure 2C). In general, connective tissue elimination and high lung epithelial progenitor enrichment is obtained after four passages, depending on the donor.

4 Quality control screening for epithelial enrichment in airway organoids by immunofluorescence (Figure 2C).
  a. During organoid passages (approximately after 4-7 passages), keep 10-20 µl of the organoid suspension (See above step 3d.). Recover intact organoids by centrifugation at 300G, 5 min at 4°C to remove the liquid Cultrex growth factor reduced BME type 2.
  b. Gently, resuspend the intact organoids in 20 µl of cold 1X PBS and transfer them to a Superfrost plus slide and incubate at RT for 10 min in a fume hood.
  c. Immediately, fix and permeabilize organoids by applying 20 µl of −30°C acetone and incubate approximately for 15 min in a fume hood until the acetone has fully evaporated. Then, use a hydrophobic pen (pap-pen) to create a hydrophobic barrier around the organoids. This will help to define the area of interest for staining.
  d. Gently rinse the organoids with 1X PBS/5% BSA/0.05% saponin.
  e. Treat organoids with Fc Receptor Block (a drop, 45 min, RT), followed by a wash step (three times with 1X PBS 5min at RT), then Background Buster treatment (a drop, 30 min, RT). One drop of each product is approximately 60 µl,
  f. Wash the organoids three times with 1X PBS (5 min at RT).
  g. Prepare the mix of primary antibodies (unconjugated, approximately 100 µl per slide, see key resources table): dilute the antibodies anti-pan-Cytokeratin (pan-CK, to reveal enrichment of epithelial cells), anti-alpha-SMA (aSMA, to reveal presence of fibroblasts) and anti-Fibronectin (FN1, to reveal presence of connective tissue) in staining buffer 1X PBS/5% BSA/0.05% saponin. Incubate in the dark for 1h at RT. Then, repeat wash step (f).
  h. Prepare the mix of secondary antibodies (Raised against the species of the primary antibody, approximately 100 µl per slide, see key resources table): dilute the antibodies anti-mouse IgG1 (specie of the anti-pan-CK antibody), anti-mouse IgG2a (specie of the anti-aSMA antibody) and anti-rabbit IgG (specie of the anti-Fibronectin antibody) in staining buffer 1X PBS/5% BSA/0.05% saponin. Incubate in the dark for 30 min at RT. Then, wash step (f).
  i. Finally, counterstain sections with 1 µg/ml final concentration (in 1X PBS, 5 min, RT) of 4’,6-diamidino-2-phenylindole (DAPI), then repeat wash step (f).
  j. Whole mount organoids slide with Fluoromount-G (Thermo Fisher Scientific) and cover glass. Then, acquire using a confocal microscope (for high resolution images). Analysis can be performed using Imaris software (Bitplane).

#### Primary human lung organoid-derived air-liquid interface (ALI) generation, differentiation & viral exposure

Timing: Approximately 6 weeks

This part describes the workflow to culture primary human lung organoid-derived air-liquid interface (ALI) cultures, along with TEER measurement to monitor the quality of tight junction formation (Figure 3).

5 Prepare the transwells (24 well plate) and then seed with lung organoid-derived single-cell suspension for ALI culture generation.
  a. Pre-coat the Transwells (24 well plate) with Collagen I (rat) (100 µl, final concentration of 30µg/ml in 1X PBS) for 1 hour at 37°C in a cell incubator.
  b. Rinse the wells with 1X PBS and leave them submerged in 1X PBS (at 37°C in cell incubator) until cell seeding (approximately 30 min to 1h).
6 ALI culture generation includes 3 steps (Figure 3A).
  a. Prepare fresh complete medial for ALI expansion supplemented with Y-27632 as described in materials and equipment section.
  b. Thaw lung organoid-derived cell suspension by placing 1 cryovial (containing 1.5×10^6^ cells/ml) into a dry bath at 37°C for 5 min.
  c. Transfer the cells into a conical 50ml tube with fresh pre-warmed ALI Expansion media (1 cryovial with 10ml media). Centrifuge at 300 G (5 min at RT) to remove the DMSO and re-suspend in ALI Expansion media (300 000 cells/ml), plate the cells (30 000 cells/100 µl/well). From one cryovial one can normally seed ∼48 inserts.
  d. (1) Expansion step: Cultivate cells submerged in expansion media until reaching 100% confluence (approximately 10-12 days depending on the donor). Change the media every day on apical (100 µl) and basal side (500 µl).
  e. (2) Differentiation step 1: Prepare fresh complete media for ALI differentiation and maintenance supplemented with Y-27632 as described in materials and equipment section. Once 100% confluence is reached, media is changed to complete media for ALI differentiation and maintenance supplemented with Y-27632 on apical and on basal side. During this step, ALI cultures establish tight junctions. TEER measurements are performed using the EVOM2 ohm meter during day 13-19 after every media changing (3 times per week). Once TEER values rise values >500 ohms.cm^2^, this indicates a healthy confluent layer (Figure 3B).
  f. (3) Differentiation step 2: Prepare fresh, complete medial for ALI expansion and complete media **without** Y-27632 for ALI differentiation and maintenance as described in materials and equipment section. Airlift ALI cultures by removal of media from the apical side while maintaining media supply to the basal side (to be changed 3 times per week). When, cultures are air-lifted, it is important that Y-27632 is no longer added to the media. This initiates differentiation into a pseudo-stratified epithelium starts. Typically, after airlift the TEER values drop to approximately 250-500 ohms.cm^2^ depending on the donor. ALI cultures should be checked under the microscope regularly for evidence of beating cilia and mucus production, this step is critical. It takes approximately 4 weeks post-airlift until full differentiation is achieved.
7 ALI culture exposure to SARS-CoV-2 virus should be performed in a biosafety level (BSL) 3 facility. <u>Troubleshooting 4</u>.
  a. Before viral exposure, remove mucus by applying pre-warmed 1X PBS apically and incubate 15 min at 37°C. Gently pipette up and down to remove mucus. Repeat this step until all mucus is removed; if the apical wash is viscous, it indicates the presence of mucus, and the wash step should be repeated. Be careful to not disrupt the cell layer.
  b. Change basal media and proceed to viral exposure by applying 10^5^ PFU virus apically (approximately 25-100 µl). Incubate the mock inserts (non-infected controls) with infection medium containing no virus with similar volume to the infected conditions.
  c. Perform viral infection in a kinetic fashion; harvest inserts every day from 1 to 6 post-infection for analysis.
  d. To assess response to virus multiple readouts can be used and some examples will be summarized in this protocol.

**Figure 3:**
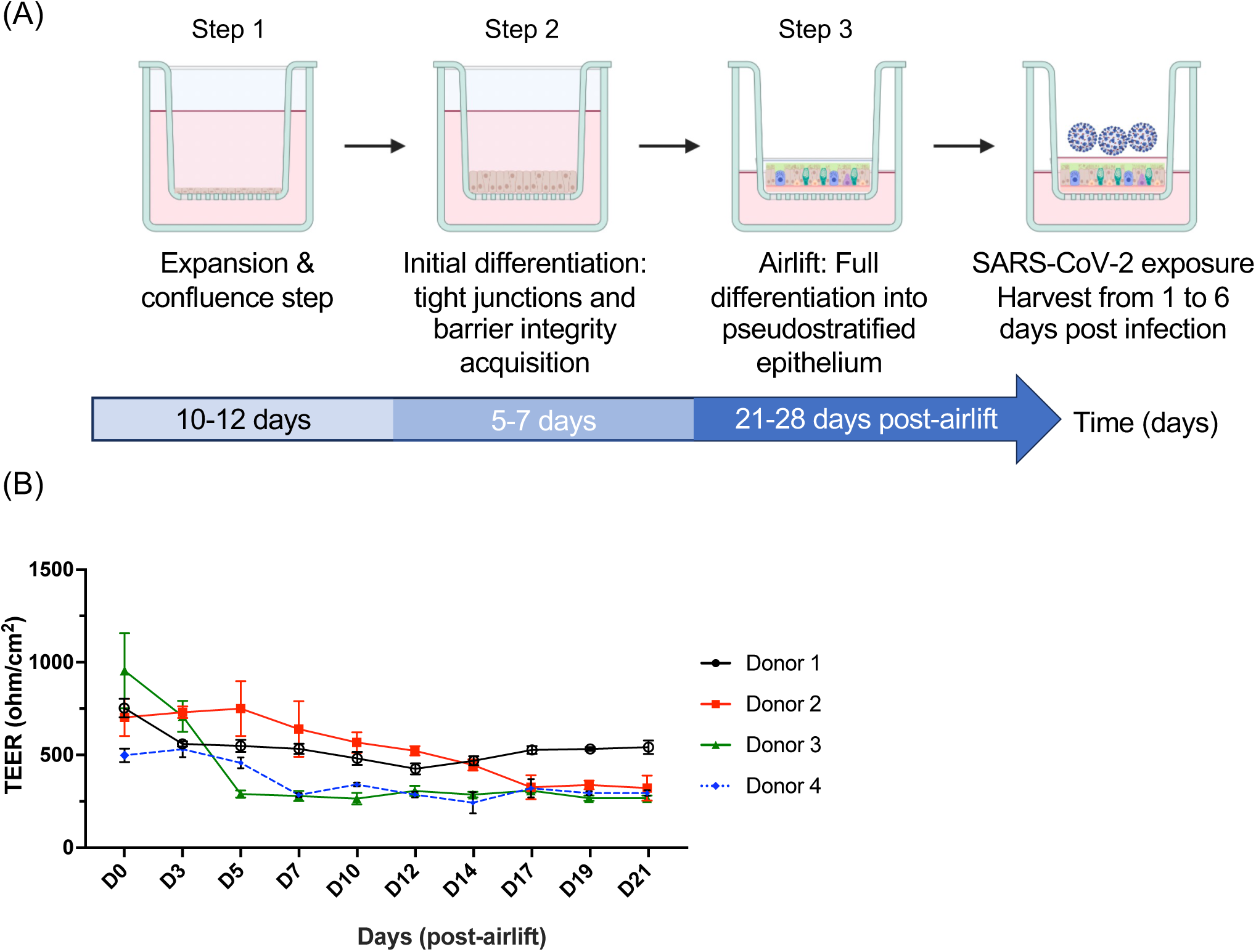
Generation of primary human lung organoid-derived ALI cultures to study response to virus: (A) Schematic experimental design for primary human lung organoid-derived ALI culture generation: (1) cell expansion in submerged culture to obtain confluence at 100%; (2) initial differentiation in submerged cultures to foster tight junctions and barrier integrity, monitored by TEER values (>500 ohms.cm2). (3) TEER goals are achieved, cultures are transitioned to airlift by removal of apical media, which initiates final differentiation into pseudo-stratified epithelia. Cultures are monitored for a minimum of 4 weeks for the presence of beating cilia and mucus production. (B) Measurement of trans-epithelial electrical resistance (TEER, ohm.cm^2^) with error bars (mean ± SD), 3 measurement per time-point performed 3 times per week starting when the epithelium is confluent, one representative experiment per donor (four donors). Fully pseudostratified differentiated ALI cultures are obtained within 3-4 weeks (post-airlift) from primary lung organoid progenitors.

**Critical:** Note that during ALI generation, the TEER evaluation is critical, if ALI is airlifted without reaching the TEER peak, the cultures will collapse. For apical infection between 25-100 µl of viral suspension could be used for viral infection. Using larger volumes may result in tissue damage and collapse of the culture.

#### Examples of readouts to assess response to viral exposure

Timing: Approximately 3 weeks

This part briefly describes five methods to assess the response to virus: Plaque assays (Viral titers); Flow cytometry (virus detection on dissociated tissue); Immunofluorescence on non-dissociated tissue (quality controls steps during ALI culture generation, cell composition, virus detection, immune marker detection); RNA data (bulk RNA) and GeoMx analysis (probe based), (Figure 4.A). After viral exposure, ALIs are harvested daily from 1 to 6 days post-infection. At least 1 insert is used for each readout at each time-point.

8 Plaque assays for viral titers on apical supernatants:
  a. Supernatant collection: Add 150 µl of pre-warmed 1X PBS to the apical side of each ALI culture and place in a cell incubator for 15 minutes. Remove supernatants by pipetting and store at −80°C in a BSL3 facility until further use.
  b. Serially dilute (10-fold) each apical supernatant from infected and non-infected ALI (mock) cultures in 1X PBS containing 1% BSA (adjust the volume for a 12 well plate).
  c. Overlay each dilution per condition on pre-seeded confluent VeroE6 cell (for USA/WA1-2020) or Vero-E6 TMPRSS2 (for Beta, Delta and Omicron) monolayers, and incubate at 37°C for 1 hour with gentle shaking every 5 min. When performing in 12-well plates, infect each well with 200 µl of diluted samples.
  d. Overlay a solution containing 2% Oxoid agarose mixed with 2X MEM supplemented with 0.3% FBS and incubate for 72h at 37°C.
  e. Fix with 4% formaldehyde solution (500 µl, overnight, room temperature). Then, wash with 1X PBS, 3 times 5 min at RT.
  f. Visualize viral infection by immune staining with SARS-CoV-2 NP antibody (1C7C7) primary antibody for 1.5 h at room temperature (RT) with gentle shaking. Then, wash with 1X PBS, 5 min at RT.
  g. Incubate with an HRP-conjugated secondary anti-mouse antibody (1:5000) for 1h at RT with gentle shaking Then, wash with 1X PBS, 5 min at RT.
  h. Reveal plaques using True Blue peroxidase substrate (Sera Care, 250 µl for a 12 well plate for around 10 min) and calculate titers as plaque-forming units per ml (PFU/ml) for every sample: 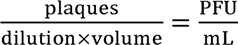
9 Flow cytometry: for intracellular viral detection on dissociated tissue
  a. Single cell generation: Wash ALI cultures with 1X PBS (150 µl on apical and 500 µl on basal side), then, treat with 0.05% trypsin on apical (150 µl) and basal chamber (500 µl) for 15 min, 37°C. Pipette up/down to dissociate the ALI tissue and transfer the cell suspension into a 15 ml tube.
  b. Neutralize trypsin with an equal volume of 1X PBS containing 2% FBS (650 µl, RT). Then, wash by centrifugation at 300 G for 5 min.
  c. Resuspend cells in a dispase I/DNase I solution (3 ml), (final concentration in 1X PBS, 1.9U/ml diaspase I (Sigma) and 0,1mg/ml DNase I (Sigma)). Incubate for 10-15 min at 37°C in a dry bath. Then, wash by centrifugation at 300 G for 5 min at RT.
  d. Viability evaluation: resuspend cells in 1X PBS and stain with zombie live dead stain (Zombie aqua or zombie green; Biolegend) in 1:350 dilution for 15 min at RT.
  e. Centrifuge at 300 G for 5 min (RT). Then, resuspend the cells in 4% methanol-free formaldehyde (100 µl/condition) and fix overnight at 4°C.
  f. Add 1ml of 1X perm wash buffer (BD Biosciences) per tube containing fixed cells and centrifuge at 300 G for 5 min (RT).
  g. Resuspend the pellet in 100ml perm wash buffer with 1:100 dilution of Alexa fluor 488-conjugated or 1:200 of Alexa fluor 647-conjugated SARS-CoV-2 N antibody (1C7C7) and incubate in dark at RT for 45 min.
  h. Add 1ml perm wash buffer and centrifuge at 300G for 5 min (temp).
  i. Finally, resuspend the pellet in 200ml 1X PBS+ 1% BSA and analyze by flow cytometry (BD, Flowjo) for live SARS-CoV-2 NP positive cells (Figure 4A).
10 Immunofluorescence: for intracellular viral detection on non-dissociated tissue
  a. Fix ALI cultures using 4% PFA (16% PFA methanol free diluted in 1X PBS, 15 min at RT), wash with 1X PBS, then maintain in 1X PBS at 4°C until further processing (stable approximately for 36 months).
  b. For each condition cut ALI mesh in half using a sterile scalpel, embed in OCT and snap freeze in liquid nitrogen, (Figure 4B).
  c. Cut frozen sections at 8 µm, air dry on Superfrost plus slides (Figure 4C) and use a hydrophobic pen (pap-pen) to create a hydrophobic barrier around the tissue sections. This will help to define the area of interest for staining.
  d. Fix sections with 4% PFA (15 min, RT), then, wash three times with 1X PBS (5 min).
  e. Permeabilize with 1X PBS/0.1% Triton-X (15 min, RT).
  f. Wash tissue sections three times with 1X PBS (5 min, RT).
  g. Treat sections with Fc Receptor Block (45 min), followed by a wash step (f), then Background Buster treatment (30 min, Innovex Bioscience). One drop of each product (approximately 60 µl), incubation at room temperature (RT). Then, repeat the wash step (f).
  h. Stain first with unconjugated antibodies, (1hr at RT), and then apply secondary antibodies (30 min, RT). Dilute antibodies in staining buffer 1X PBS/5% BSA/0.05% saponin. Between the unconjugated and secondary antibodies incubation repeat the wash step (f).
  i. Repeat wash step (f) and add mouse serum to saturate secondary antibodies. Dilute normal mouse serum 1/20 in 1X PBS and incubate for 15 minutes at room temperature. Then, repeat the wash step (f).
  j. Continue the staining with directly conjugated antibody mix for 1 hour at room temperature and then repeat wash step (f).
  k. Depending on the staining panel (See below, Panel 3), stain for actin filaments with 10 nmol units/ml final concentration (in 1XPBS, 15 min, RT) of Phalloidin ATTO647N (Sigma 65906), then wash step (f).
  l. Finally, counterstain sections with 1 µg/ml final concentration (in 1XPBS, 5 min, RT) of 4’,6-diamidino-2-phenylindole (DAPI), then wash step (f).
  m. Mount ALI slides with Fluoromount-G (Thermo Fisher Scientific) and cover glass. Acquire using a confocal microscope (for high resolution images) or a widefield microscope (for histocytometry). Analysis can be performed using Imaris software (Bitplane).

**Figure 4:**
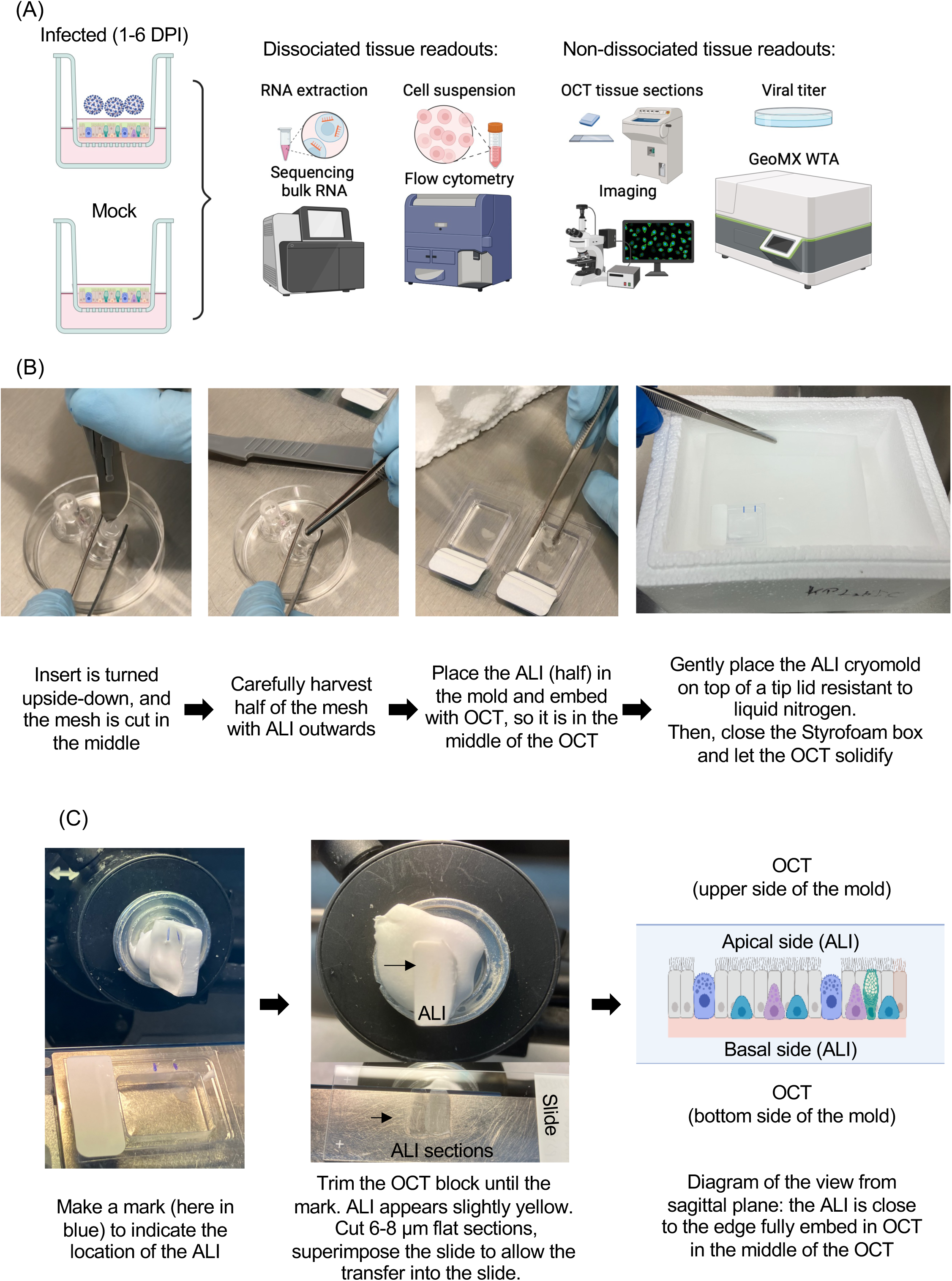
Readouts to assess response to virus, ALI culture OCT embedding and OCT cutting. (A) Schematic workflow presenting five methods to assess response to virus. Flow cytometry and RNA extraction require tissue dissociation versus Immunofluorescence, plaque assay (viral titer on apical supernatant) and GeoMx WTA do not require tissue dissociation. (B) Procedure of embedding ALIs in OCT and cryopreservation. First, turn the insert upside-down, and with a scalpel cut the mesh in the middle and partially around the edges, enough to hold the mesh and pull-out from the insert. Be careful not to damage the cell layer. Place the ALI mesh on top of a layer of OCT and cover it with OCT to fully embed the ALI insert. Gently, with a pipette tip make sure the ALI is not curved or too close to the bottom or surface of the cryomold. Then, snap freeze in liquid nitrogen. To ensure no liquid nitrogen (LN) enters the tissue, the cryomold should be placed on top of a plastic lid resistant to LN. (C) ALI section cutting and transfer to Superfrost plus slides. Make a mark with a sharpy pen to indicate the location of the ALI culture on the cryomold then on the solidify OCT block. This tip will facilitate the OCT trimming until the mark and then visualize the ALI (slightly in yellow). The diagram allows to picture the configuration of the ALI culture in the OCT block. Finally, ALI culture sections could be transferred into Superfrost plus slides. These steps will ensure good quality sections for further experiments. **Method video S1: Video illustrating the procedure of embedding ALI cultures in OCT.**

In the context of our study, three panels of antibodies are detailed here as examples (See Key resources table):

| Panel 1 | Panel 2 | Panel 3 |
| --- | --- | --- |
| Cell composition:<br>See Figure 5A | CSF3 and CCL20 expression<br>on infected ALI: See Figure 5B | Infected and non infected<br>ALI: See Figure 5C |
| Unconjugated Abs:<br>Anti-SCGB1A1<br>Anti-Acetyl alpha-tubulin | Unconjugated Abs:<br>Anti-CCL20 | Unconjugated Abs: NA |
| Secondary Abs:<br>Anti-rat IgG AF488<br>Anti-mouse IgG2b AF555 | Secondary Abs:<br>Anti-rabbit IgG AF555 | Secondary Abs: NA |
| Serum saturation: Yes | Serum saturation: Yes | Serum saturation: NA |
| Conjugated Abs:<br>Anti-cytokeratin 5 AF594<br>Anti-MUC5AC AF700 | Conjugated Abs:<br>Anti-CSF3 AF594<br>Anti-NP SARS-CoV-2 AF488<br>(1C7C7) | Conjugated Abs:<br>Anti-NP SARS-CoV-2<br>AF488 (1C7C7) |
| Phalloidin ATTO647N: NA | Phalloidin ATTO647N: NA | Phalloidin ATTO647N:<br>Yes |

Note: it is important that the ALI culture is correctly embedded in OCT (Figure 4B-C and Method video S1). These steps will ensure the quality of sections for immunofluorescence and GeoMx WAT.

11 Bulk RNA sequencing:
  a. First dissociate ALI as described in steps (9.a.) to (9.c.). From 1 insert approximately 1-2 x 10^6^ cells are normally recovered. After cell suspension is washed by centrifugation (300 G, 5min, RT) then, isolate total RNA using Direct-zol RNA MicroPrep kits (zymo research, around 200 µl of Direct-zol buffer) following the manufacturer’s protocols.
  b. Perform a DNase treatment using the RNase-free DNase set (Qiagen) according to the manufacturer’s protocols.
  c. Prepare cDNA libraries using polyA mRNA selection and Kapa Stranded mRNA-Seq Library Prep kit (Kapa Biosystems) according to the manufacturer’s instructions. Perform paired-end sequencing (i.e., 2 × 150 bp) of stranded total RNA libraries using Illumina Novaseq or similar.
  d. Use FastQC for quality control ^10^, remove rRNA contamination with MultiQC^8^ and trim reads with BBDuk tool^11^.
  e. Map viral reads to the FDA-ARGOS SARS-CoV-2 reference sequences (FDAARGOS_983, with bowtie^7,12^, and map human reads to GRCh38 genome using bowtie2 (https://www.nature.com/articles/nmeth.1923). Gene expression is quantified using RSEM^6^. Batch correction is applied using R package SVAseq^13^. Finally, use DESeq2^9^ for read normalization and differential gene expression.

Note: it is important that time-matched non-infected conditions are added to the experimental design. This will ensure the quality and accuracy of RNA-Seq data analysis. As an example of resulting data see Figure 6 A, which is a Heatmap representing differentially expressed genes (DEGs) over the time in response to SARS-CoV-2 variants. ALI from four donors were infected with SARS-CoV-2 and harvested for sequencing at 1, 2, 3, 4, 5 and 6 dpi and mock-infected (control media without virus) samples collected from days 1 to 6 as well.

12 GeoMx whole Transcriptome Atlas: this assay, is a probe-based method of reporting *in situ* RNA data analysis that can detect expression of 18,676 genes. It includes a first step with immunofluorescence (3 targets + nuclei: basal cells (cytokeratin 5), virus NP and Spike SARS-CoV2 proteins) to define the regions of interest (ROI).
  a. Cut 6 µm thick sections from ALI culture-OCT blocks, and air dry on Superfrost plus slides. Slides can be stored in a secured −80 °C freezer until use.
  b. Prepare PFA (4% in 1X PBS) fixed-frozen tissue slides according to the GeoMx NGS automated Leica Bond RNA Slide Preparation Manual (NanoString, MAN-10131-03). For more details, please refers to Nanostring website.
  c. Load slides into the slide holder of the GeoMx digital spatial profiling (DSP) instrument and cover with 2 mL of manufacturer buffer S.
  d. Stain each slide with the following antibodies and scan using example exposure times: respectively for the morphology markers nuclei Syto83 (Cy3/568nm, 60ms), Cytokeratin 5 (CK5, FITC/525nm, 200ms), SARS-CoV-2 Spike (Texas Red/615nm, 250ms), and SARS-CoV-2 Nucleocapsid (Cy5/666nm, 200ms).
  e. Select regions of interest (ROIs) based on the antibody staining. See Figure 6B, representative images of ALI infected at 1, 2 and 6 day-post infection (DPI) with ROIs selection. The thin white polygons represent the ROI strategy selection for the apical cytokeratin 5-cells (CK5-) vs basal side CK5+.
  f. Expose selected ROIs to UV photocleavable barcode RNA probes. Then, process DSP and PCR according to the manufacturer’s protocol. Sequence purified libraries using an Illumina NovaSeq6000 or similar machine.
  g. GeoMX data processing and QCs according to Nanostring protocols.
  h. Perform data analysis in R (v4.2.0). Produce graphics using ggplot2 unless otherwise stated (v3.3.6).
  i. Differentially expressed genes: Use a Wilcoxon rank-sum test to identify DEGs between different regions of interest. Adjust p values using Benjamini-Hochberg multiple test correction. Use p value cut-off of 0.05 and a logFC of 2. Identify genes variability across the whole dataset by a Kruskal-Wallis test (p < 0.05) in order to show clustering of different ROI groups by heatmap.
  j. Gene expression heatmaps: Produce Heatmaps using the ComplexHeatmap package (v2.12.1). Heatmaps use by-row scaling, and ROIs are first grouped by infection type, ordered by Day, and then clustered using the default hierarchical clustering algorithm.

**Figure 6:**
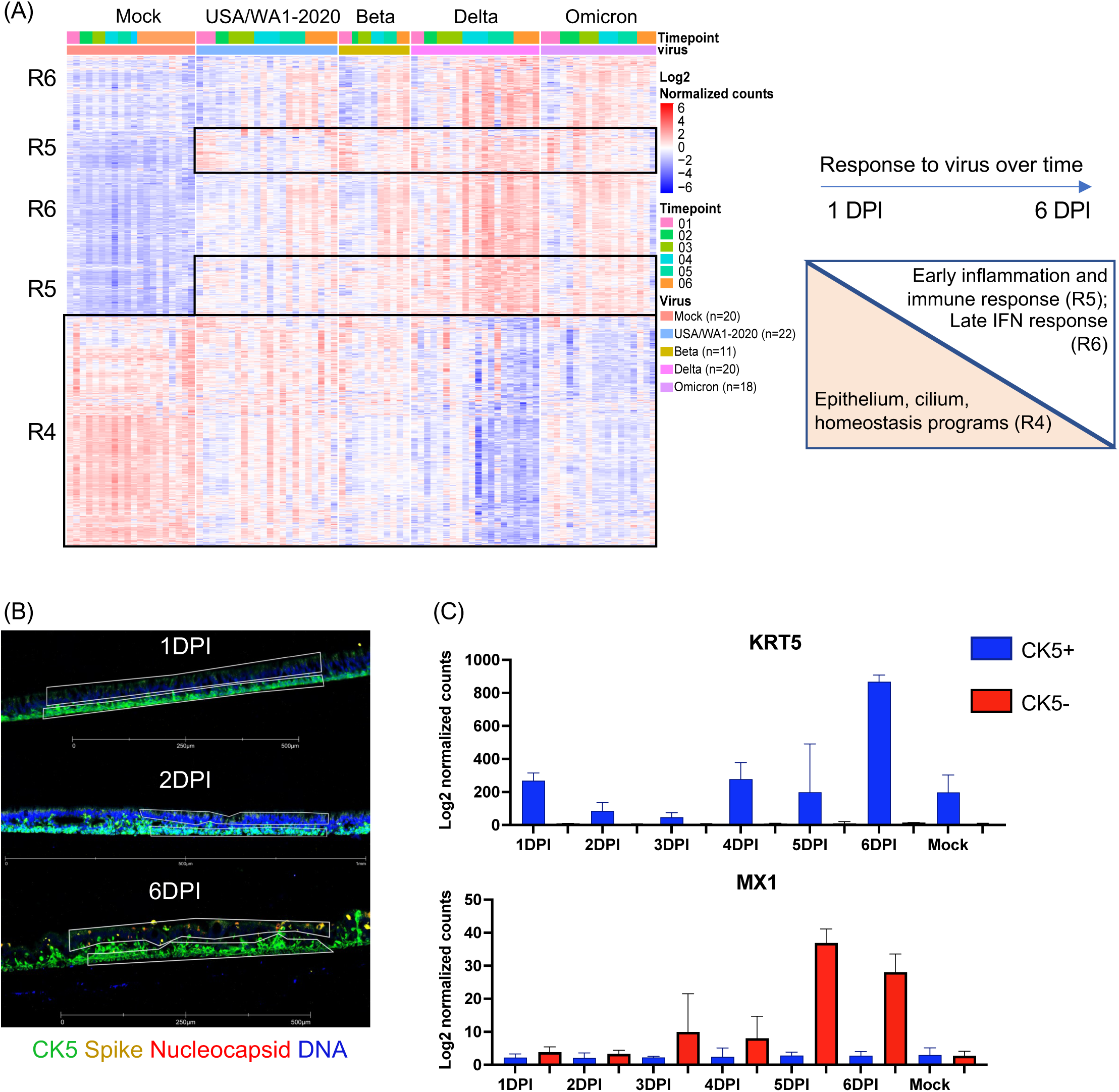
Examples of expected outcomes of transcriptional response to SARS-CoV-2 variants and ROI selection for GeoMx data. (A) Heatmap representing differentially expressed genes over the time in response to SARS-CoV-2 variants. ALI cultures from four donors were infected with SARS-CoV-2 and harvested for sequencing at 1, 2, 3, 4, 5 and 6 dpi and mock-infected (control media without virus) samples collected from days 1 through 6 days as well. The sequencing was performed in multiple batches with at least 2 independent experiments at each time-point, the cut-off used for defining differentially expressed genes: |logFC| > 1; adjusted p-value < 0.01; normalized counts > 10. Rows represent individual transcripts and columns represent individual biological replicates ordered by timepoints and SARS-CoV-2 variants. Batch effect was removed using SVAseq R package. Three clusters were identified: Cluster R4 was enriched for cilia and epithelium maintenance signatures. Interestingly, this signature was persistently downregulated in response to Delta and Omicron variants. The cluster R5 and R6 showed a strong up-regulation of inflammatory and immune signatures for all variants starting at 1 DPI. All variants induced expression of genes associated to viral response at later time-points. This response to virus from 1 to 6 DPI is also depicted by the schematic on the right, with cluster R4 decreasing from 1 to 6 DPI whereas clusters R5 and R6 are increasing. (B) ROI selection for GeoMx data: Representative images of infected ALI culture sections with SARS-CoV-2 (10^5^PFU), at 1 DPI, 2 DPI, 6DPI stained for nuclei (Dapi), viral nucleoprotein (red), spike viral protein (yellow) and cytokeratin 5 (CK5). The thin white polygons represent selected ROIs for the apical cytokeratin 5^-^ cells (CK5^-^) vs basal side CK5^+^. Scale bar 500 μm for 1 and 6 DPI; and 1 mm for 2 DPI (white). (C) GeoMx data: Bar graphs were generated using Graphpad (Prism5) and illustrate the gene expression (log 2 normalized counts) over time within selected ROIs based on cytokeratin 5 protein expression (CK5 +) through KRT5 gene expression and based on cytokeratin 5 negative expression (CK5 -) of infected cells positive for SARS-CoV-2 Spike and NP, through MX1 as part of the anti-viral response which is mainly increased at later time-points.

**Critical:** GeoMx was initially optimized for FFPE tissue but has been validated on PFA fixed-frozen fixed tissues. Staining must be validated for other antibodies.

### Expected outcomes

The generation of primary human lung organoids has become an invaluable and widely utilized technique in biomedical research. Their use as progenitors for air-liquid interface (ALI) cultures has been particularly critical in advancing our understanding of the host response to respiratory viruses such as SARS-CoV-2. By using this protocol, researchers should be able generate a large stock of organoid derived lung epithelial progenitor cells that can be used for ALI culture production. Here we also briefly describe some readouts that could be used to assess ALI development and to measure the response to virus. Notably, a simple control under brightfield microscope allows one to monitor the proper development of ALI cultures by following mucus formation and cilia beating as illustrated in Video S1. Furthermore, Figure 5A, 5B and 5C illustrate results of immunofluorescence readout to monitor respectively ALI differentiation by following cell composition, viral infection with CSF3 localization on the basal side and CCL20 on the apical side and a simple staining with anti-NP and phalloidin report levels of ALI culture infection. Flow cytometry, another protein-based method allows one to measure viral infectivity as illustrated by Figure 5D. Finally, Figure 6A, illustrates the transcriptional response to virus across time-points and across variants. Furthermore, Figure 6B-C, illustrates the quality of the sections, and the precision of ROI selection resulting at RNA level (probe-based) with a “transcriptional” separation between cells expressing cytokeratin 5 (CK5+) and those infected not expressing CK5 (CK5-) based on MX1 expression as downstream of infection.

**Figure 5:**
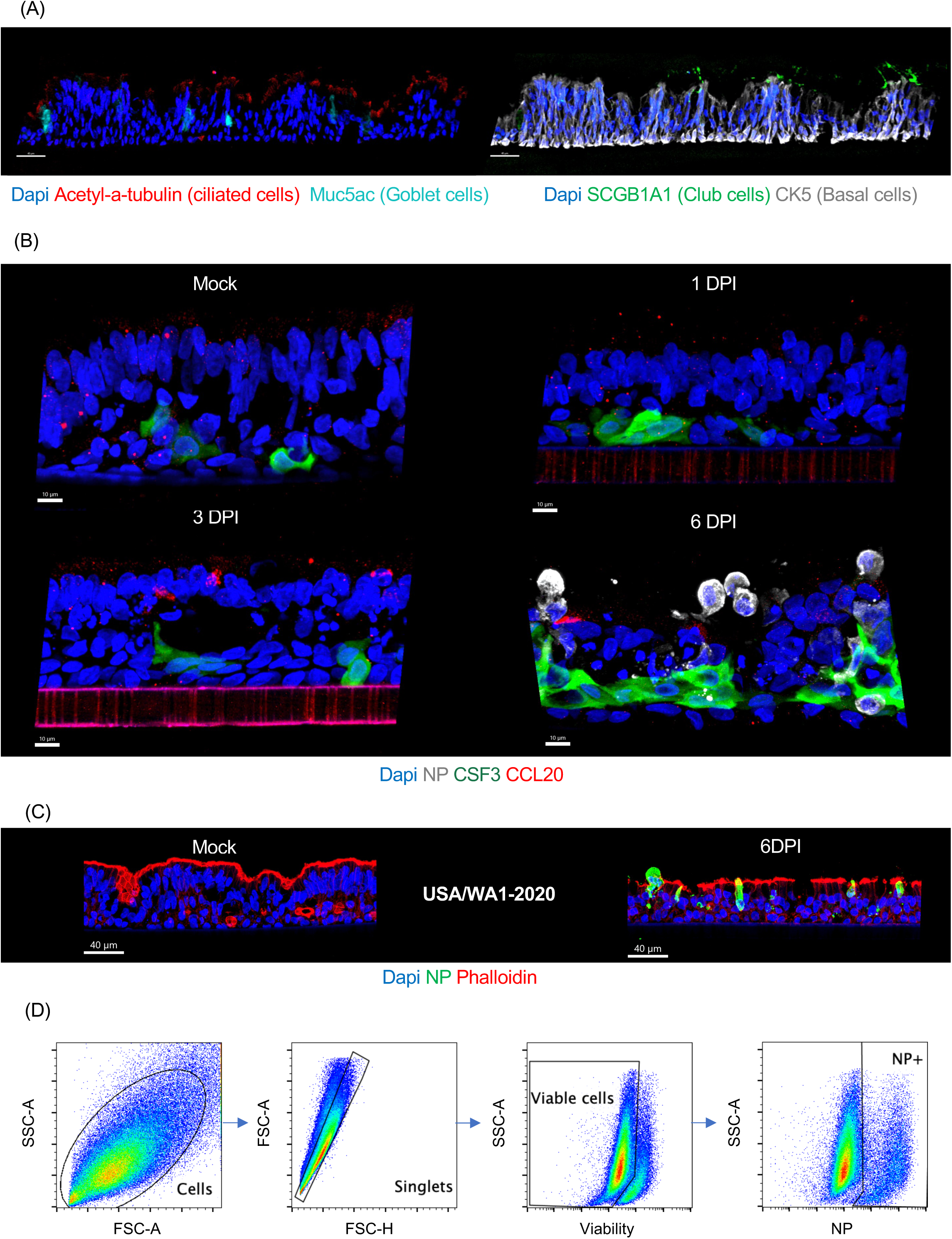
Examples of expected outcomes of immunofluorescent images and flow cytometry gating strategy. (A) Representative immunofluorescent (IF) section (8 µm) of differentiated lung organoid-derived ALI cultures. Left panel, merged figures, showing markers for ciliated cells (acetylated a-tubulin, red) and goblet cells (MUC5AC, cyan). Right panel, merged figures, showing club cells (SCGB1A1, green), basal cells (CK5, white). Nuclei (Dapi, blue). Scale bar 40 µm, in white on the left corner. (B) Representative images of ALI cultures mock-infected at 6 days and ALI infected with SARS-CoV-2 USA/WA1-2020 (105PFU) at 1, 3 and 6 days-post infection (DPI) stained for nuclei (Dapi, blue), viral NP (white) to reveal the effective viral replication, CSF3 (green) and CCL20 (red). Scale bars 10 μm, in white on the left corner. (C) Representative images of donor 3 of mock-infected (control media without virus at 6 days) and infected ALI cultures with SARS-CoV-2 (10^5^PFU) at 6 days post infection (DPI) stained for nuclei (Dapi), viral nucleoprotein (NP, green) to reveal the effective viral replication and phalloidin (Actin filament, red) to reveal tissue structure. Scale bar 40 μm, in white on the left corner. (D) Gating strategy example for flow cytometry analysis. A single cell suspension was prepared from SARS-CoV-2 infected ALI cultures. Cell suspensions were stained for viability and viral infection by using an anti-NP antibody specific to SARS-CoV-2. Plots from left to right show serial gating to identify percentages of infected (NP positive) viable cells. **Video S1: Video illustrating movement of beating cilia in ALI cultures at D34 post-airlift.**

### Limitations

This protocol involves the use of viable frozen lung tissue from whole lung donors to generate primary human lung organoids, which are then used to generate air-liquid-interface (ALI) cultures for studying the response to viruses. However, due donor-to-donor variations in lung tissue, there may be differences observed during the expansion of organoids and variation in ALI culture composition and response to virus. Additionally, primary cell cultures are susceptible to microbial and fungal outgrowth during organoid and ALI generation process. To

prevent cross-contamination between wells or plates and maintain the integrity of the cultures, the following measures are taken: Prepare one tube of media per plate, and to minimize the risk of cross-contamination, change tips between each well to avoid transferring potential contaminants. Contaminated plates should be immediately bleached and discarded to prevent further spread of contaminants. By implementing these precautions, researchers can reduce the risk of contamination and maintain the quality and consistency of primary human lung organoid cultures, allowing for more reliable and reproducible experimental results.

### Troubleshooting

#### Problem 1

Incomplete or partial tissue digestion, related to the section “Human lung tissue viable freeze thawing & lung airway organoid generation” step (2 d) involving collagenase digestion.

#### Potential solution

- Transfer the tissue into a petri dish and with sterile slides mechanically shred the tissue to break the connective tissue.
- Centrifuge at 300G for 5min (temp) and repeat the step (d) with new 10 ml of complete media for primary lung airway organoids (AO) containing 1–2 mg ml collagenase I on an orbital shaker at 37°C for an additional hour.

#### Problem 2

Incomplete red blood cell lysis, related to “Human lung tissue viable freeze thawing & lung airway organoid generation” step (2 f).

#### Potential solution

- Make sure cell culture grade water is used to dilute the red blood cell lysis buffer and not PBS.
- Incubate samples on ice for 3-4 min and then stop the reaction by the addition of 3ml of 1X PBS.

#### Problem 3

Loss of cells during organoid passaging due to their high adherence to the plastic, related to “Human lung tissue viable freeze thawing & lung airway organoid generation” step (3).

#### Potential solution

- For every step involving organoids, tips, pipettes, tubes, are pre-coated with 1X PBS with 1% BSA to avoid organoid adherence.

#### Problem 4

Loss of cells during the viral infection, due to the excess of apical buffer and/or mucus. Related to “Primary human lung organoid-derived air-liquid interface (ALI) generation, differentiation & viral exposure“ step (7).

#### Potential solution

- Prevent mucus buildup on the apical side by washing every 48-72 hours before infection.
- For every infection, use between 25-100ul virus infection medium. More than 100ul will likely drown the cells, if left for more than 24-48 hours.
- If using more than 100ul, the infection medium should be removed after 3 hours of infection.

## Resource availability

### Lead contact

Further information and requests for resources and reagents should be directed to and will be fulfilled by the lead contact: Karolina Palucka or Adam Williams.

### Materials availability

This study did not generate new unique reagents.

### Data and code availability

- All sequencing data generated mentioned here has been deposited to GEO and is publicly available: <u>GSE225603</u>
- This paper does not report original code.
- Any additional information required to reanalyze the data reported in this work paper is available from the lead contact upon request.

## Supporting information

Method Video S1 link to Figure 4

## Acknowledgments

We thank our tissue donors. The Microscopy, Single Cell Biology, Genome Technology, and CTRS Scientific Services of The Jackson Laboratory. We thank Randy Albrecht for support with the BSL3 facility and procedures at the Icahn School of Medicine at Mount Sinai (ISMMS).

## Author contributions

Conceptualization, D.C.C., and K.P.; Methodology, A.W., F.M, D.C.C., and K.P.; Validation, D.C.C., and S.J.; Formal analysis and Data Curation, M.Y.; Investigation, D.C.C, S.J., M.S., J.M., M.C. (Megan Callender), M.C., A.C., J.G.-B.D., T.-C.W., and F.M.; Resources, A.G.-S., and M.S.; Writing-Original Draft, D.C.C.; Writing-Review & Editing, D.C.C., S.J., A.W., M.Y., M.S., A.G.-S., and K.P.; Funding acquisitions, A.W., A.G.-S. M.S. and K.P.; Supervision, D.C., A.G.-S., M.S., A.W., and K.P.

## Declaration of interests

The A.G.-S. laboratory has received research support from GSK, Pfizer, Senhwa Biosciences, Kenall Manufacturing, Blade Therapeutics, Avimex, Johnson & Johnson, Dynavax, 7Hills Pharma, Pharmamar, ImmunityBio, Accurius, Nanocomposix, Hexamer, N-fold LLC, Model Medicines, Atea Pharma, Applied Biological Laboratories and Merck, outside of the reported work. A.G.-S. has consulting agreements for the following companies involving cash and/or stock: Castlevax, Amovir, Vivaldi Biosciences, Contrafect, 7Hills Pharma, Avimex, Pagoda, Accurius, Esperovax, Farmak, Applied Biological Laboratories, Pharmamar, CureLab Oncology, CureLab Veterinary, Synairgen, Paratus and Pfizer, outside of the reported work. A.G.-S. has been an invited speaker in meeting events organized by Seqirus, Janssen, Abbott and Astrazeneca. A.G.-S. is inventor on patents and patent applications on the use of antivirals and vaccines for the treatment and prevention of virus infections and cancer, owned by the Icahn School of Medicine at Mount Sinai, New York, outside of the reported work.

The M.S. laboratory has received unrelated funding support in sponsored research agreements from Phio Pharmaceuticals, 7Hills Pharma, ArgenX and Moderna. K.P. is a stockholder in CueBiopharma and Guardian Bio, scientific advisor to Cue Biopharma and Guardian Bio and co-founder of Guardian Bio. K.P. declares unrelated funding support from Guardian Bio (current) and MERCK (past). All additional authors declare no competing interests.

